# Development and patterning of a highly versatile visual system in spiders

**DOI:** 10.1101/2023.12.22.572789

**Authors:** Luis Baudouin Gonzalez, Anna Schönauer, Amber Harper, Saad Arif, Daniel J. Leite, Philip O. M. Steinhoff, Matthias Pechmann, Valeriia Telizhenko, Atal Pande, Carolin Kosiol, Alistair P. McGregor, Lauren Sumner-Rooney

**Affiliations:** Oxford University Museum of Natural History, University of Oxford, Parks Road, Oxford OX1 3PW, UK; Department of Biological and Biomedical Sciences, Oxford Brookes University, Gipsy Lane, Oxford OX3 0BP, UK; Enara Bio, Science Park, Bellhouse Building Level 3, Sanders Rd, Littlemore, Oxford OX4 4GA, UK; Department of Biosciences, Durham University, Stockton Road, Durham DH1 3LE, UK; Zoologisches Institut und Museum, Universität Greifswald, Loitzer Strasse 26, 17489 Greifswald, Germany; Department of Developmental Biology, Universität zu Köln, Zuelpicher Strasse 47B, 50674 Köln, Germany; School of Biology, St Andrews University, St Andrews KY16 9ST, UK; Leibniz Institute for Biodiversity and Evolution, Museum für Naturkunde, Invalidenstrasse 43, 10115 Berlin, Germany

## Abstract

Visual systems provide a key interface between organisms and their surroundings, and have evolved in many forms to perform diverse functions across the animal kingdom. Spiders exhibit a range of visual abilities and ecologies, the diversity of which is underpinned by a highly versatile, modular visual system architecture. This typically includes eight eyes of two developmentally distinct types, but the number, size, location, and function of the eyes can vary dramatically between lineages. Previous studies of visual system development in spiders have confirmed that many components of the retinal determination gene (RDG) network are conserved with other arthropods, but so far, comparative studies among spiders are lacking. We characterised visual system development in eight species of spiders representing a range of morphologies, visual ecologies, and phylogenetic positions, to determine how these diverse configurations are formed, and how they might evolve. Combining synchrotron radiation tomography, transcriptomics, in situ hybridisation, and selection analyses, we characterise the repertoires and expression of key RDGs in relation to adult morphology. We identify key molecular players, timepoints, and developmental events that may contribute to adult diversity, in particular the molecular and developmental underpinnings of eye size, number, position, and identity across spiders.

## Introduction

Vision is a key evolutionary innovation that changed the course of animal history more than half a billion years ago. Eyes and visual systems have since evolved dozens of times across the animal tree of life, in a wide variety of forms suited to their respective needs. Such needs can range from the simple detection of shadows to the resolution of fine spatial, temporal, or spectral details for visual communication, and eyes themselves can vary from small pigmented spots to large, sophisticated, and regionalised structures. The configuration of the whole visual system is also highly variable: although many taxa have one pair of lateral, cephalic eyes, others exhibit more complex architectures comprising more eyes, potentially performing different functions in parallel (Buschbeck and Bok 2023). For example, many insects have two or three ocelli in addition to their compound eyes, while molluscs such as scallops and chitons may have hundreds of duplicated eyes spread across the body. Different configurations of eyes presumably offer different advantages, yet the evolution of visual system architectures remains a somewhat underexplored area.

Spiders provide an excellent opportunity to study how and why visual system architectures diversify. Most spiders have four pairs of eyes of two distinct types, providing the basis for a highly versatile modular system (Figure 1). This is demonstrated by the substantial variation of several key factors including eye size, number, position, and function (Morehouse et al. 2017; Morehouse 2020). For example, families such as Caponiidae, Tetrablemmidae, and Dysderidae can have between two and six eyes of only one type, jumping spiders (Salticidae) and ogre-faced spiders (Deinopidae) exhibit dramatic functional and structural divergence between their four eye pairs, and the eyes of some male erigonines are placed on elaborate modifications of the carapace. This morphological and functional variation enables a rich diversity of visual ecologies and behaviours among spiders, notably including functional tetrachromatic vision, the use of celestial polarisation compasses, complex visual communication, and sophisticated stalking and prey capture hunting modes (e.g. Dacke et al. 1999; Zurek et al. 2015; Jakob et al. 2018).

**Figure 1.**
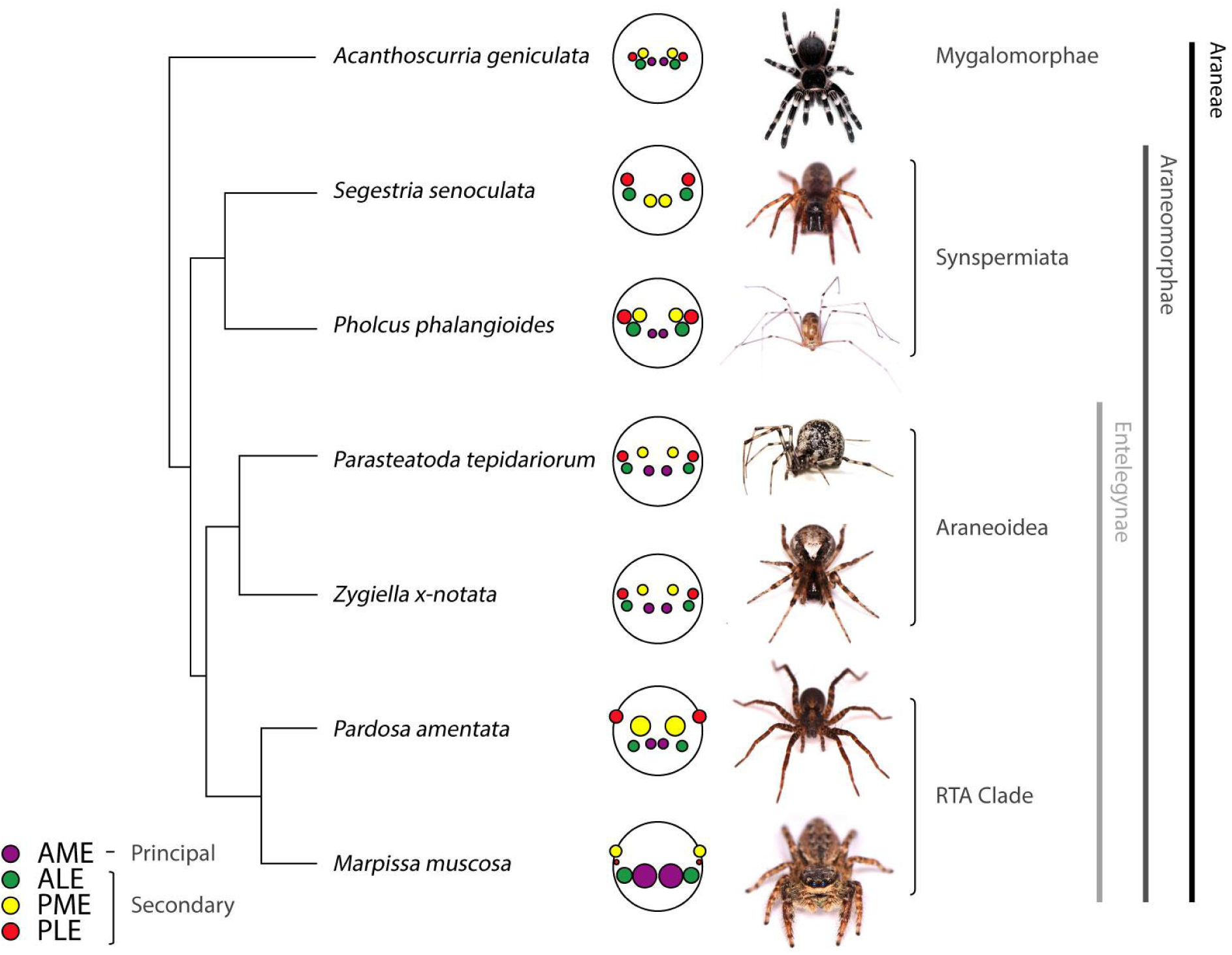
Species were selected for this study on the basis of their phylogenetic distribution and morphological disparity in adult visual systems. We examined visual system development in *A. geniculata* (Theraphosidae), *S. senoculata* (Segestriidae), *P. phalangioides* (Pholcidae), *P. tepidariorum* (Theridiidae), *Z. x-notata* (Araneidae), *P. amentata* (Lycosidae), and *M. muscosa* (Salticidae), representing four major clades of spiders and a range of visual ecologies and hunting modes. Spiders have up to four pairs of eyes - one pair of principal (anterior median) eyes, and three pairs of secondary eyes. Note that, in line with observations made in this study, the posterior-most pair of eyes in *M. muscosa* is designated the posterior median eyes. Images kindly provided by Sam J. England (*S. senoculata, P. phalangioides, Z. x-notata, P. amentata* and *M. muscosa*), Grace Blakeley (*P. tepidariorum*), and Matthias Pechmann (*A. geniculata*). Topology and branch lengths based on divergence time estimates from Shao and Li (2018).

Despite this diversity, spider visual systems are united by a conserved anatomical blueprint and clear homologies, even between lineages separated by hundreds of millions of years of evolution. They are therefore well suited for the comparative study of visual system evolution (Morehouse et al. 2017).

Morphological diversification is underpinned by changes in developmental processes between lineages. This can include changes in the regulation and expression of genes in the underlying gene regulatory networks. Visual system development is regulated by a network of retinal determination genes (RDGs), which appear to be highly conserved across large phylogenetic distances. For example, transcription factors including PAX6, Otd, and Sine oculis play key roles in the development of eyes in taxa as diverse as vertebrates, insects, and molluscs (Arendt 2003).

However, we know relatively little about visual system development and how it has evolved to generate morphological and functional diversity in spider visual systems. The eyes form from two pairs of primordia: one appearing above the anterior furrow, giving rise to the principal eyes, and the other at the lateral extremes of the head, which eventually splits and invaginates to form the three secondary eye pairs (Homann 1971; Samadi et al. 2015; Schomburg et al. 2015). Studies of the model species *Parasteatoda tepidariorum* and *Cupiennius salei* have characterised key components of the retinal determination pathways and their expression in developing embryos (Samadi et al. 2015; Schomburg et al. 2015; Baudouin-Gonzalez et al. 2022). These recovered many orthologs of the RDG network described in detail in *Drosophila melanogaster* (Gaspar et al. 2019; Chen and Desplan 2020), including many genes present in duplicate, likely as a result of the ancient whole-genome duplication in arachnopulmonates (Schwager et al. 2017). These studies demonstrated the expression of these RDGs in the developing eye primordia and indicated support for the homology of principal eyes with insect ocelli, and of secondary eyes with insect compound eyes, based on similarities in the expression of genes such as *otd/ocelliless* and *Six3/Optix.* They also highlighted a possible pathway to determining eye identity, as each eye pair expressed different combinations of RDGs. However, a number of striking differences emerge between even these two species: for example, *otd2* expression was detected in the anterior median eyes (AMEs) of *P. tepidariorum* but not *C. salei*, and vice versa for *Pax6.2* (Samadi et al. 2015; Schomburg et al. 2015; Baudouin Gonzalez et al. 2022). In *C. salei*, *Pax6.2* expression was also detected relatively late in eye development; given that it is implicated in the initiation of eye development in a wide range of other organisms (Gehring, Walter; Ikeo 1999; Hartmann et al. 2003; Callaerts et al. 2006; Gehring 2014), this implies that the structure and regulation of RDG networks is not necessarily conserved, even where similar suites of genes are expressed. Furthermore, spatiotemporal expression patterns indicate that while the role of Wnt signalling, which suppresses the expression of RDGs in *D. melanogaster*, is likely to be conserved in *P. tepidariorum*, this does not appear to be the case for *hedgehog* or *decapentaplegic* (Baudouin-Gonzalez et al. 2022).

Studies of eye development in *P. tepidariorum* and *C. salei* to date hint at several possible mechanisms that could facilitate the diversification of spider visual systems: distinct developmental origins of the two eye types, the retention of duplicate RDGs, the expression of different RDG combinations in different eye pairs, and different spatiotemporal expression patterns of the same RDGs, could all contribute to both phylogenetic divergence between taxa and developmental divergence between eye pairs. However, a wider comparative approach, using a broader phylogenetic and morphological range, is essential to identify which of these vary consistently between taxa, and which correlate with morphological and functional variation.

We characterised RDG repertoires and spatiotemporal expression patterns in eight spider species representing the four major clades of extant spiders (Mygalomorphae, Synspermiata, Araneoidea, and the RTA clade) and a range of adult morphologies and visual ecologies to explore possible developmental mechanisms underpinning key characteristics of visual system configuration: eye size, number, location, and function. We conducted selection analyses for RDGs to explore the potential divergence between lineages and duplicated genes, and tracked the gross development of the eye primordia and the central nervous system (CNS) in four key species using synchrotron radiation tomography to align this with the observed RDG expression patterns.

While RDG repertoires were highly conserved across all species examined, we identified important differences in their spatial and temporal expression patterns that provide insight to the patterning of adult diversity. Notable features of the visual system, including eye size, number, and arrangement, are reflected in the expression patterns of selected genes relatively early in development, some of which also indicate signs of subfunctionalisation between putative ohnologs.

## Materials and methods

### Spider culture and embryo collection

Adult spiders of *Marpissa muscosa* (Greifswald, Germany), *Pardosa amentata* (Oxford and Nottinghamshire, UK, and Berlin, Germany), *Pholcus phalangioides* (Oxford, UK), *Segestria senoculata* (Oxford, UK) and *Zygiella x-notata* (Oxford, UK) were collected and kept at 25°C with a 12:12h light:dark cycle for cocoon collection. *Parasteatoda tepidariorum* embryos were collected from an in-house culture at Oxford Brookes, kept under the same conditions, and *Acanthoscurria geniculata* embryos were collected from an in-house culture at the Universität zu Köln (Pechmann 2020). An egg sac of *Drassodes* sp. was collected from a drystone wall in West Yorkshire, UK. Embryos were staged using Mittmann and Wolff (2012) as a guide and fixed for *in situ* hybridisation (ISH) following the protocol in Akiyama-Oda et al. (2016), with minor modifications as described in Baudouin-Gonzalez et al. (2021).

### Synchrotron radiation tomography

Staged embryos were fixed in 70% ethanol and mounted in pipette tips for scanning at the TOMCAT beamline of the Swiss Light Source (Paul Scherrer Institute) (Stampanoni et al. 2006). Samples were scanned using a monochromatic beam (energy 16 keV), with propagation distances between 10-100 mm, and combined magnifications of 4x (effective voxel size 1.65 μm, LuAG:Ce 100 μm scintillator), 10x (effective voxel size 0.65 μm, LuAG:Ce 20 μm scintillator), or 20x (effective voxel size 0.325 μm, LuAG:Ce 20 μm scintillator). Samples were rotated through 180°, across which 2000 projections were recorded with exposure times of 70 ms (4x), 120 ms (10x), or 200 ms (20x). In-house bespoke software was used to reconstruct slices from projections, with Paganin filtering (delta = 1e-7, beta = 3e-9) (Paganin et al. 2002) and depth reduction to 8-bit tiffs (Marone and Stampanoni 2012). Data were cropped in Fiji (Schindelin et al. 2012) and models were produced in Amira v.2021 (Thermo Fisher), using volume rendering for the outer surfaces and manual segmentation for the central nervous system.

### RNA extraction and transcriptome assembly

Total RNA was extracted per species from mixed stage pooled embryos of *Z. x-notata*, *S. senoculata* and *Drassodes* sp using QIAzol lysis reagent following the standard protocol (Qiagen). Libraries were prepared using a TruSeq RNA kit (including polyA selection) and sequenced on the NovaSeq platform (100 bp PE, Edinburgh Genomics). Erroneous k-mers were removed using rCorrector v.1.0.4 (default settings) (Song and Florea 2015) and uncorrectable read pairs were discarded using a custom Python script (available at https://github.com/harvardinformatics/TranscriptomeAssemblyTools/blob/master/FilterUncorrectabledPEfastq.py courtesy of Adam Freeman). Adapter sequences were identified and removed, and low-quality ends trimmed using TrimGalore! v.0.6.5 (phred score cutoff =5) (Capella-Gutiérrez et al. 2009). Quality assessment was performed between each step using FastQC v.0.11.9 (Andrews 2010). The resulting processed reads were used to perform *de novo* transcriptome assembly with Trinity v.2.11.0 (default settings) (Haas et al. 2013). Transcriptome completeness was verified with Busco v.5.0.0 (Seppey et al. 2019) using only the longest isoform per gene (default settings, arachnida_obd10 lineage).

### Identification and phylogenetic analysis of RDGs

RDGs were identified using tBLASTn (e-value 0.05; Camacho et al. 2009), using protein sequences previously identified in *P. tepidariorum* (Schomburg et al. 2015; Baudouin-Gonzalez et al. 2022) as queries against the available transcriptomes of *A. geniculata* (PRJNA588224), *P. phalangioides* (Turetzek, Torres-Oliva, Kaufholz, Prpic, Posnien, unpublished data), *M. muscosa* (PRJNA707377), *P. amentata* (PRJNA707377), *Charinus acosta* (PRJNA707377), and *Euphrynichus bacillifer* (PRJNA707377), and the newly assembled transcriptomes of *Z. x-notata*, *S. senoculata*, and *Drassodes* sp.. Protein sequences were predicted from the identified transcript sequences using the ORFfinder NCBI online tool (https://www.ncbi.nlm.nih.gov/orffinder/; the “any sense codon” setting was used to retrieve protein sequences from fragmented transcripts).

To confirm gene identity and orthology, we performed phylogenetic analysis using the full-length protein sequences of RDGs from 12 spiders (*P. tepidariorum, Z. x-notata, Araneus ventricosus, Argiope bruennichi, Stegodyphus dumicula, C. salei, P. amentata, M. muscosa, Drassodes* sp*, S. senoculata, P. phalangioides* and *A. geniculata*), two amblypygids (*C. acosta* and *E. bacillifer*), a scorpion (*Centruroides sculpturatus*), a tick (*Ixodes scapularis*) and two insects (*D. melanogaster* and *Tribolium castaneum*). RDG protein sequences from *P. tepidariorum* and *C. salei* were retrieved from previous studies (Samadi et al. 2015; Schomburg et al. 2015; Baudouin-Gonzalez et al. 2022). RDG protein sequences from *A. ventricosus, A. bruennichi, S. dumicula, C. sculpturatus, I. scapularis, T. castaneum* and *D. melanogaster* were retrieved from their respective NCBI proteomes. Alignments were performed in Clustal Omega using the default parameters (https://www.ebi.ac.uk/Tools/msa/clustalo/). Maximum likelihood phylogenetic analysis was performed in RAxML-NG v.1.0.2 (Stamakis 2019), using the best fitting model as suggested by ModelTest-NG v.0.1.7 (JTT+I+G4+F for *atonal*, JTT+I+G4+F for *otd*, VT+I+G4+F for Pax genes, JTT+I+G4+F for Six genes, PMB+I+G4+F for *dac* and JTT+I+G4+F for *eya*) and the automatic bootstrapping algorithm. Trees were visualised using FigTree v.1.4.4 (http://tree.bio.ed.ac.uk/software/figtree).

### Cloning and probe synthesis

cDNA synthesis was carried out using the QuantiTect reverse transcription kit (Qiagen). Gene fragments were amplified from cDNA by PCR reaction and cloned into the pCR®4-TOPO®TA vector using the TOPO®TA cloning kit for sequencing (ThermoFisher Scientific). A list of primers used is provided in Supplementary Table 1. RNA probes were synthesized using T7 (10881775001, Roche) or T3 RNA polymerase (11031163001, Roche), depending on the orientation of the cloned fragment in the pCR®TOPO®TA plasmid, using DIG RNA labelling mix (11277073910, Roche).

### In situ hybridisation

Whole-mount ISH was performed following the protocol of Prpic et al. (2008), with minor modifications as described in Baudouin-Gonzalez et al. (2021). Ethanol treatment was used to reduce background staining as described in Baudouin-Gonzalez et al. (2021). Embryos were counterstained with DAPI (1:2000; 10236276001, Roche) for ∼20 min and stored in PBS-T at 4°C. Imaging was performed using a Zeiss Axio Zoom V.16 and DAPI overlays were processed in Photoshop CS6 (Adobe). Embryos between stages 9.1 and 13.2 were used, with ISH performed at each of stages 9.1/9.2, 10, 11, 12, 13.1, and 13.2 when availability allowed.

### Selection analyses

For selection analyses we also queried the genes and transcripts from the above species against genomes and genome annotations generated in Leite et al. (in prep). Blastn (Camacho et al. 2009) was used to identify putative sequences (e-value 0.05) and sequences were extracted with samtools (Danecek et al. 2021) (transcriptomes and genome annotations) or bedtools getfasta (Quinlan and Hall 2010) (genomes). Hits from genome annotations were collapsed based on the gene ID to remove isoforms, whereas hits from transcriptomes and genomes were collapsed using cd-hit (Li and Godzik 2006) with a cluster threshold of 90% and cd-hit-lap to cluster overlapping sequences. Where multiple hits were recovered against a single gene, nucleotide alignments were inspected to eliminate non-overlapping potential gene fragments; the longest fragments that overlapped with the most conserved regions were used for phylogenetic analysis.

Coding sequences for each gene were then retrieved and translated into amino acid sequences. To preserve the reading frame, protein sequences were aligned using the L-INS-i method implemented in MAFFT 7 (Katoh and Standley 2013). Protein alignments were then used to build corresponding alignments of nucleotide coding sequences in the PAL2NAL software (Suyama et al. 2006). Regions that are predominantly gaps or are hard to align were removed with the Gblocks (Castresana 2000).

Phylogenetic trees for selection analyses were constructed for each gene using IQ-TREE (Nguyen et al. 2015) with automatic selection of the best substitution model (Kalyaanamoorthy et al. 2017). Node support was evaluated with the ultrafast bootstrap method (Hoang et al. 2018).

We used the CodeML program in the PAML 4 (Yang 2007) package to characterise the action of natural selection on coding sequences. The ratio of rates of nonsynonymous to synonymous substitutions (dN/dS or ω), was calculated and compared for different scenarios. An ω value below 1 indicates purifying selection, ω=1 characterises neutral evolution, and ω>1 suggests positive selection.

First, we used the branch model (Nielsen and Yang 1998; Yang 1998), which assumes different ω ratio parameters for different lineages of the hypothesised phylogeny to evaluate selection pressure on different sets of gene copies of spiders. For this, paralogous spider species groups were labelled foregrounds and tested sequentially for each gene. The significance of the obtained results was assessed using the likelihood ratio test (LRT) with null model M0, which assumes the same ω for all branches. Next, we performed branch tests on separate lineages selected as foreground branches due to additional duplications or distinctive expression patterns.

The branch-site model was then applied to detect potential amino acid sites under positive selection along these lineages. The Bayes empirical Bayes method (Yang et al. 2005) was used to identify codon sites potentially under positive selection on the foreground branches. Statistical significance was evaluated using LRT and twice the log-likelihood difference between null and alternative models, compared to a 50:50 mixture of a chi-square distribution with one degree of freedom and a point-mass at 0 (Zhang et al. 2005).

## Results

### Gross development of the eyes and CNS

As previously described by Schomburg et al. (2015) and Baudouin-Gonzalez et al. (2022) for *P. tepidariorum* and Samadi et al. (2015) for *C. salei*, as well as by Homann (1971) for several other species, the principal and secondary eyes form from separate primordia. The principal eyes originate in the non-neurogenic ectoderm at the anterior-most tip of the head lobes, around or anterior to the anterior furrow, and subsequently migrate posteriorly and ventrally during head closure. These are visible as distinct *sine oculis* (*so*) expression patterns from around stage 9.2. The secondary eyes arise from the ventro-lateral part of the head lobes, starting from a single pair of primordia that divide into three between stages 10.2 and 12. These form as small indentations of the epithelium (Figure 2) that invaginate and become covered by the developing lens, as described by Homann (1971). These invaginations are visible from stage 13.1 onwards in synchrotron scans of most species. The secondary eye pairs are named according to their developmental homology across species, based on previous descriptions of their division and migration in *P. tepidariorum*: the dorsal-most pair migrates into the centre of the head during head closure to form the PMEs, the ventral-median pair forms the ALEs, and the mid-lateral pair form the PLEs (Schomburg et al. 2015). Note that in *M. muscosa*, the eye pair that is highly reduced in adults shares developmental homology with the PLEs of all other species, but has historically been described as the PMEs. The eye pair previously described as the salticid PLE is homologous to the PMEs in other species. We use this new nomenclature hereafter, in line with the developmental homology. This also aligns with the designation of the PMEs and PLEs in *P. amentata* and other lycosids, where the size difference between the secondary eye pairs is similarly pronounced.

**Figure 2.**
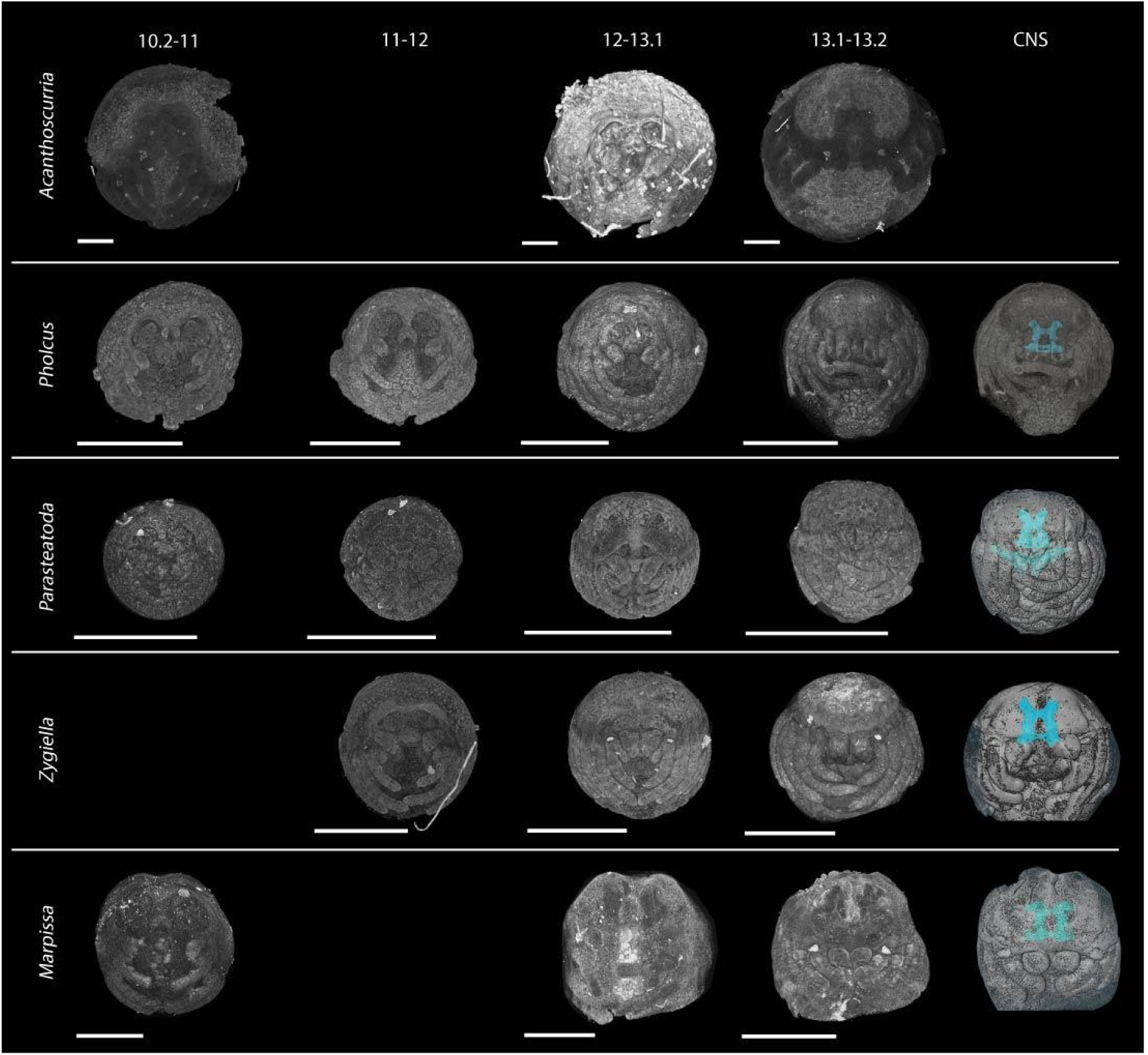
Gross development of spider embryos and their visual and central nervous systems. Models reconstructed from synchrotron scans of ethanol-preserved embryos (see methods). The central nervous system (blue) was visible by stage 13.1-13.2 in all species except *Acanthoscurria geniculata.* Black arrowheads indicate the position of the principal eyes, red arrowheads indicate the position of the secondary eyes. All scale bars are 500 μm.

The developing CNS is visible in synchrotron scans from around stage 12-13. By stage 13.1, the circumoesophageal ring formed by the connection of the syncerebrum (‘supraoesophageal ganglion’) to the suboesophageal ganglion is clearly visible, with the exception of *A. geniculata* (Figure 2). Also visible are two prominent nerve bundles projecting anteriorly from the protocerebrum, which may contribute to the developing optic neuropils. However, connections to the eye primordia were not visible even by stages 13.2 and 14.

### RDG repertoires are mostly conserved in spiders

RDG repertoires have been previously described in two spider species, *P. tepidariorum* and *C. salei* (Samadi et al. 2015; Schomburg et al. 2015).

To assess whether spider RDG repertoires have diverged in gene content, we identified orthologs of *so*, *Pax6*, *Otd*, *Six3*, *dac*, *eya* and *ato* for seven additional species, using available transcriptome data as well as new transcriptome data acquired for the purpose of this study (Supplementary Table 2). Overall, spider RDG repertoires are well conserved, with little variation of gene copy number (Figure 3). Similar to previous findings in *P. tepidariorum* and *C. salei* (Samadi et al. 2015; Schomburg et al. 2015), we identified two copies of each RDG in most species surveyed, the only exception being *eya*, for which a single copy was always retrieved (Figure 3). However, we did detect gene copy number variation in some species (Figure 3).

**Figure 3.**
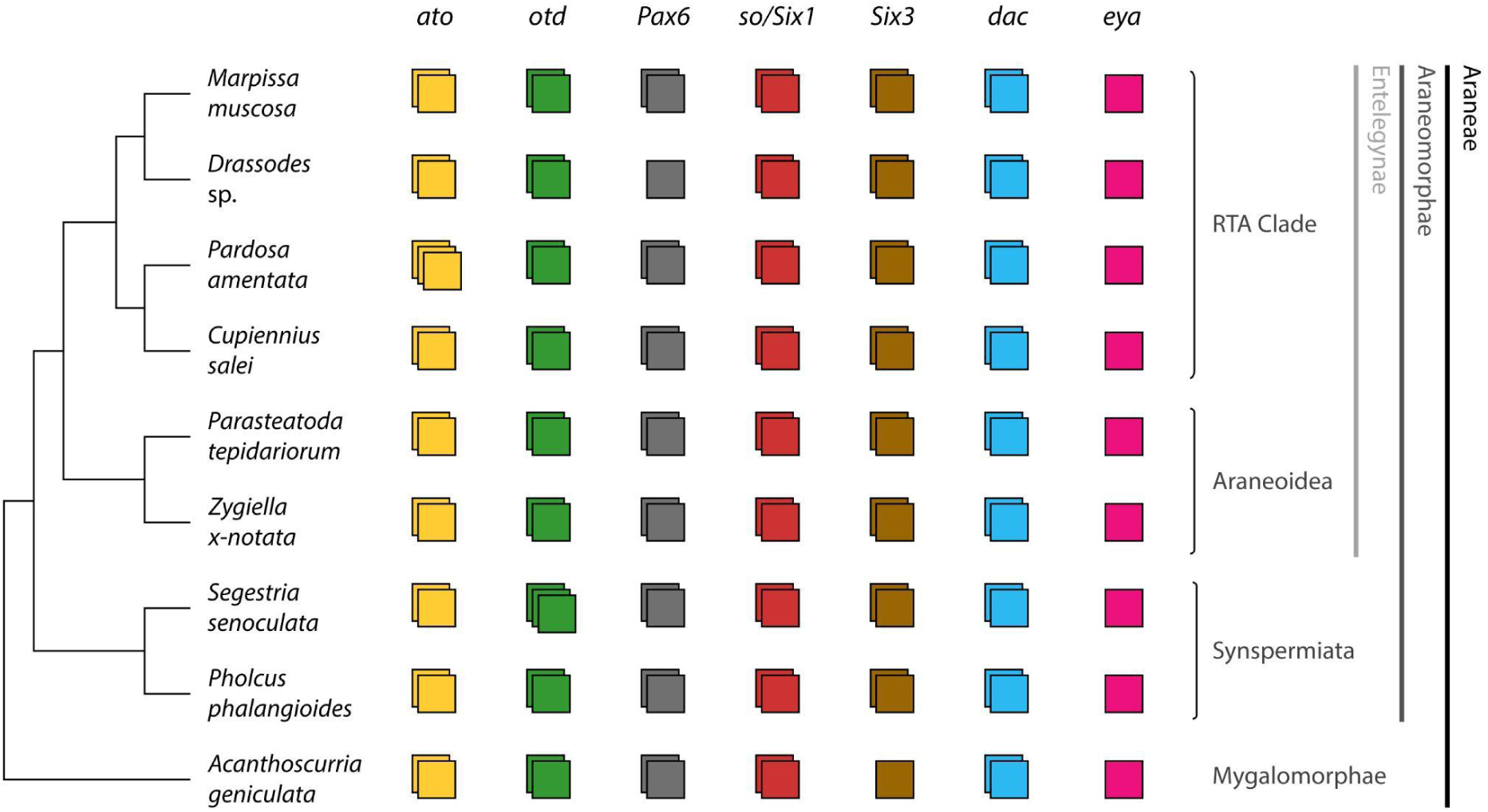
Retinal determination gene repertoires in spiders. Most species examined exhibited conserved RDG repertoires, with duplicates likely retained from an ancient whole-genome duplication in arachnopulmonates. Notable exceptions included *eyes absent*, which was present in single-copy in all species, and an additional duplicate of *orthodenticle* in *Segestria senoculata* and *atonal* in *Pardosa amentata*.

*P. amentata* has an additional copy of *ato* compared to the other species surveyed, with two of the three copies, *Pa-ato2* and *Pa-ato3*, showing high sequence conservation at the DNA level, possibly representing a very recent lineage-specific duplication (Supplementary File 1, Supplementary Figure 1). We also identified a third copy of *otd* in *S. senoculata*; two of these copies, *Ss-otd1.1* and *Ss-otd1.2*, are orthologous to *Pt-otd1*, and *Ss-otd2* is orthologous to *Pt-otd2* (Figure 3) (Supplementary File 3, Supplementary Figure 3). Interestingly, in *P. phalangioides*, the species most closely related to *S. senoculata* in our analysis, both copies of *otd* appear to be orthologous to *Pt-otd1*, forming a well-supported clade with *Ss-otd1.1* and *Ss-otd1.2* (Supplementary Figure 3). Therefore, it appears that an ortholog of *Pt-otd2* was lost in *P. phalangioides*, with *Pp-otd1.1* and *Pp-otd1.2* originating from a duplication specific to Synspermiata. Lastly, only one copy of *Pax6* was found in the transcriptome of *Drassodes* sp., orthologous to *Pt-Pax6.1* (Supplementary File 4, Supplementary Figure 4), and a single copy of *Six3* was present in the transcriptome of *A. geniculata*, which appears to be orthologous to *Pt-Six3.2* (Supplementary File 5, Supplementary Figure 5). Whether these are truly absent from the genomes of these animals, rather than undetected in the transcriptomes, cannot be confirmed at this time.

### Expression of sine oculis orthologs

Two copies of *so* were previously reported in *P. tepidariorum*, *Pt-so1* and *Pt-so2* (Schomburg et al. 2015). Our analysis of their expression patterns was consistent with previous descriptions by Schomburg et al. (2015). *Pt-so1* expression was detected in all developing eye pairs. A pair of expression domains at the anterior edge of the head at stage 10.2 correspond to the primary eye primordia, which migrate towards the final position of the AMEs by stage 13.2 following head closure (Figure 4G-G”, black arrowheads). A second pair of expression domains appear at stage 10.2 in the lateral edge of the head; these correspond to the secondary eye primordia, which then split into three distinct domains at stage 12 to form the ALEs, PMEs, and PLEs (Figure 4G-G”, red arrowheads). *Pt-so2* expression was only detected in the ALEs (Figure 4H-H”).

**Figure 4.**
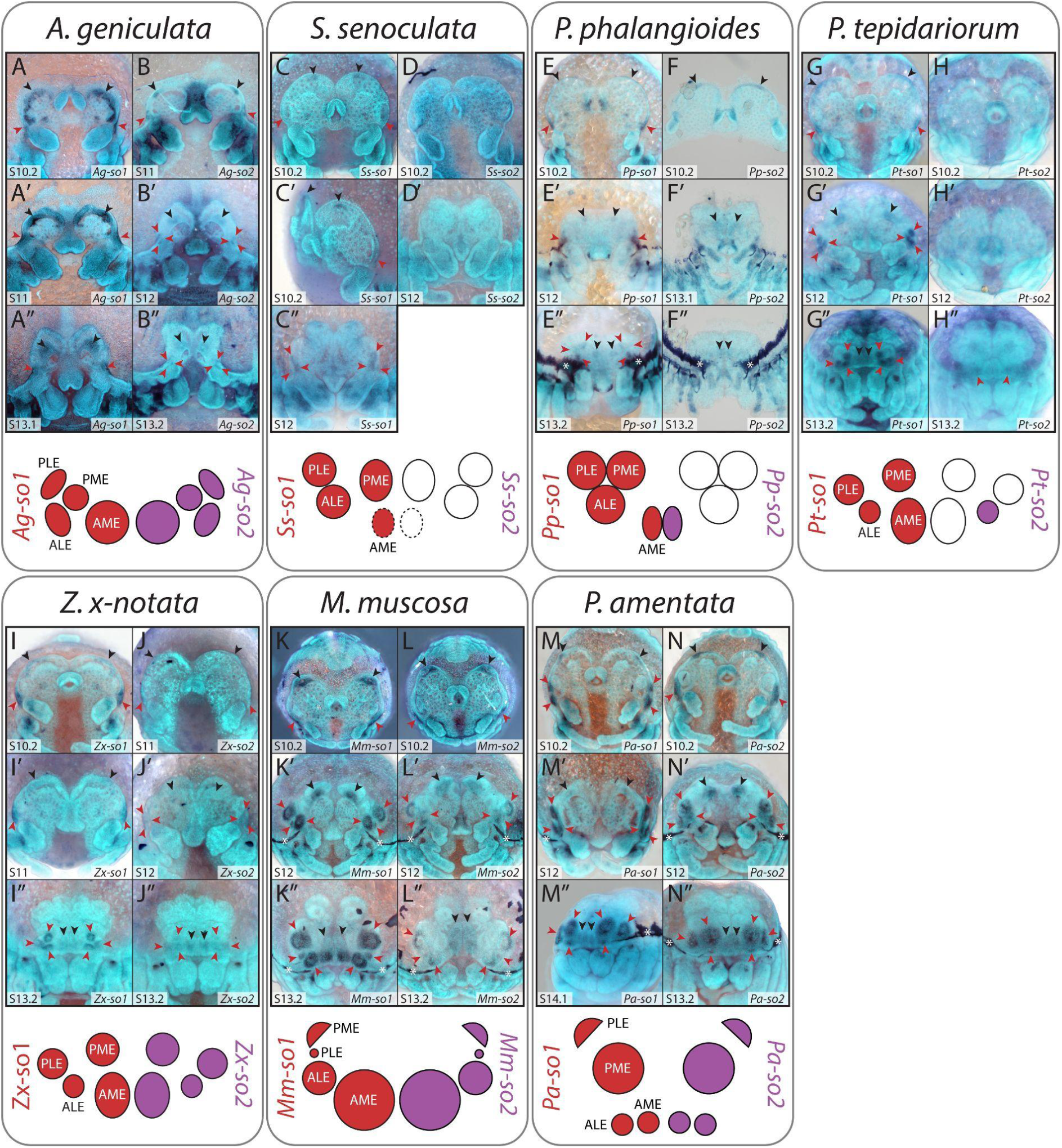
Expression of *sine oculis* orthologs in developing spider embryos. Two copies of *so* were recovered from each species. Orthologs of *so1* were expressed in all developing eyes, in all species examined, from stage 10.2 onwards. Expression of *so2* orthologs was detected from stage 10.2 in all eye primordia in *Acanthoscurria geniculata, Zygiella x-notata, Marpissa muscosa,* and *Pardosa amentata*, as well as in the AMEs of *Pholcus phalangioides* and the ALEs of *Parasteatoda tepidariorum*. Black arrowheads indicate principal eye primordia, red arrowheads indicate secondary eye primordia, and asterisks indicate artifactual staining of the developing cuticle.

For all species analysed, at least one copy of *so* is expressed in all eye primordia (Figure 4), similar to *Pt-so1* in *P. tepidariorum* (Figure 4G-G”), but the expression of *Pt-so2* orthologs is less consistent among species.

*Ag-so1* has broader expression domains than any ortholog in other species: they extend beyond the putative developing eyes, and expression in the secondary eye primordia does not split cleanly into three distinct domains at stage 13.1 (Figure 4A-A”). *Ag-so2* expression is more restricted to the eye primordia and expression in the secondary eye primordia splits into three domains starting at stage 12 (Figure 4B-B”).

In both *P. phalangioides* and *S. senoculata*, the expression domains of *Pp-so1*, *Pp-so2* and *Ss-so1* in the developing AMEs appear to be relatively smaller than in the other analysed species (Figure 4C-F”, black arrowheads). Intriguingly, *Ss-so1* expression is clearly visible in the region of the principal eye primordia at stage 10.2 despite the AMEs being completely absent in this family, but this expression can no longer be detected by stage 12 (Figure 4C-C”). Like *Pt-so2*, *Pp-so2* and *Ss-so2* are the only other orthologs of this gene whose expression was not detected in all eye primordia; however, unlike *Pt-so2* expression in the ALEs, *Pp-so2* is only expressed in the AMEs (Figure 4F-F”) and *Ss-so2* does not appear to be expressed in any of the eye primordia (Figure 4D and D’).

In *Z. x-notata*, *Zx-so1* has an identical expression pattern to *Pt-so1* (Figure 4G-G” and 4I-I”) The early expression domain of *Zx-so1* appears to be larger than *Zx-so2* and it becomes restricted to the edges of the secondary eyes during stage 13. *Zx-so1* expression is more intense than *Zx-so2* expression in these eye pairs.

In *M. muscosa* and *P. amentata* expression of both copies of *so* reflect differences in adult eye size. In *M. muscosa* the expression domains in the AMEs are larger than in other species analysed, and expression in the PLEs (historically described as PMEs) is greatly reduced in size (Figure 4K-L”). The expression domains of *Mm-so1* and *Mm-so2* are also generally much larger in the ALEs and PMEs (historically described as PLEs) than their ortholog’s expression in other species. Similarly, the region of expression of *Pa-so1* and *Pa-so2* is very large in the PMEs and PLEs, and comparatively smaller in the AMEs and ALEs (Figure 4M-N”). In these two species, spatial differences within *so* expression domains, particularly in the larger eye pairs, are visible from stages 12 onwards. In *M. muscosa*, *Mm-so1* is more strongly expressed in a ventral region of the AMEs at stage 13.2 (Figure 4K”), whereas *Mm-so2* expression is less intense and more uniform across the AMEs (Figure 4L”). In the secondary eyes, expression of *Mm-so1* is most intense around the eye perimeter, with a possible additional dot at the centre of the ALEs and PMEs (figure 4K’ and K’’). *Mm-so2* expression in the ALEs and PMEs appears to be restricted to a subset of cells around the distal edges of the eyes, forming a crescent-like pattern (Figure 4L’ and L”). In *P. amentata*, the region of *Pa-so2* expression is more intense and possibly larger than *Pa-so1* expression in the AMEs (Figure 4M-N’). In the secondary eyes, the expression of *Pa-so1* is very pronounced at the periphery of the PMEs and PLEs, with an additional ventral dot of intense expression, while *Pa-so2* expression forms a less pronounced crescent shape in these eye pairs and a central/ventral dot (Figure 4M’-N’’).

### Expression of orthodenticle orthologs

In *P. tepidariorum*, *Pt-otd1* is not expressed in the eye primordia but is visible in the developing brain at stage 10.2, becoming fainter and more obscure during head closure (Figure 5G-G”). Expression of *Pt-otd2* was detected in the primary eye primordia and in putative neural tissue from stage 10.2, which also becomes restricted to the midline during head closure (Figure 5H-H”). These patterns, which were already described in *P. tepidariorum* by Schomburg et al. (2015), are conserved in most species analysed (Figure 5). In *P. phalangioides*, *Pp-otd1.2* expression was faintly visible at the margin of the principal eye primordia (Figure 5F”), but the depth of this staining suggests that it is within the developing brain; this is consistent with our phylogenetic analysis, which suggests both copies are orthologous to *Pt-otd1* (Supplementary Figure 3). The *Ss-otd1.2* probe produced no clear staining in any embryonic tissue and is not depicted in Figure 5. In *S. senoculata*, *Ss-otd2* expression is clearly visible in the region of the primary eye primordia at stage 10.2 (Figure 5D), but is later restricted to the developing brain (Figure 5D’). The expression of *Mm-otd2* in the principal eye primordia is distinctly enlarged and intense compared to the expression of orthologs in other species from as early as stage 10.2 (Figure 5L-L’’), reflecting the enlargement of the AMEs.

**Figure 5.**
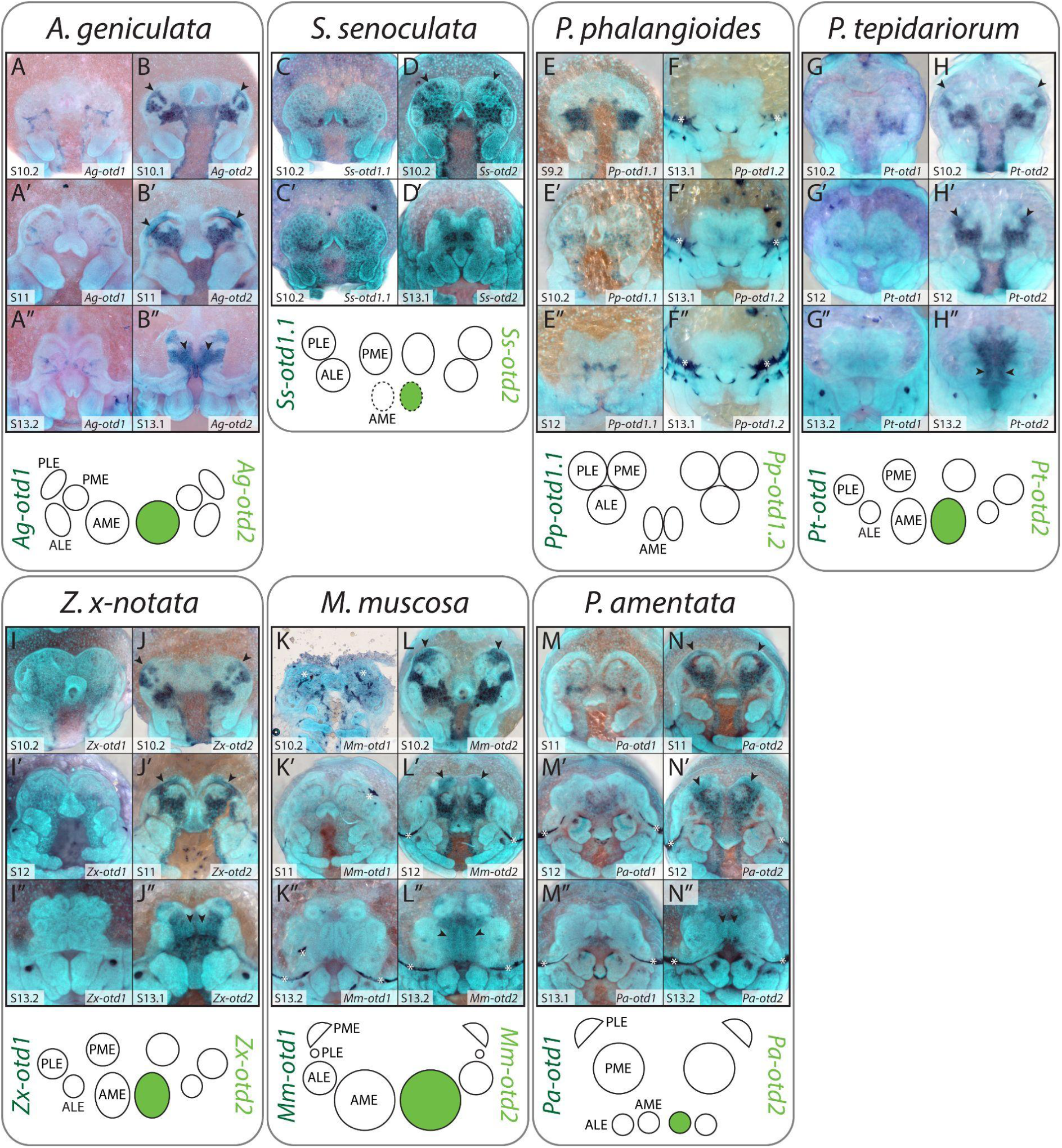
Expression of *otd* orthologs in developing spider embryos. Two copies of *otd* were recovered from each species except *Segestria senoculata*, where three copies were found. *otd1* orthologs were not expressed in any developing eyes (including *Ss-otd1.2*, not pictured). *otd2* expression was detected in the AMEs of all species except *Pholcus phalangioides*, starting from stage 10.2-11. Black arrowheads indicate principal eye primordia, red arrowheads indicate secondary eye primordia, and asterisks indicate artifactual staining of the developing cuticle.

### Expression of *Six3* orthologs

Previous descriptions of *Six3* expression in *P. tepidariorum* reported no *Pt-Six3.1* expression in any of the eye primordia, while *Pt-Six3.2* expression was detected in all secondary eye primordia, starting at stage 12, which we corroborated (Schomburg et al. 2015; Figure 6F-G”). However, we also observed two small expression domains of *Pt-Six3.1* at the edge of the non-neurogenic ectoderm from stage 10.2 and subsequently more posteriorly as the non-neurogenic ectoderm grows over the neurogenic ectoderm (Figure 6F-F”, black arrowheads). At stage 12 its location is similar to that of the *Pt-so1* expression in the AMEs.

**Figure 6.**
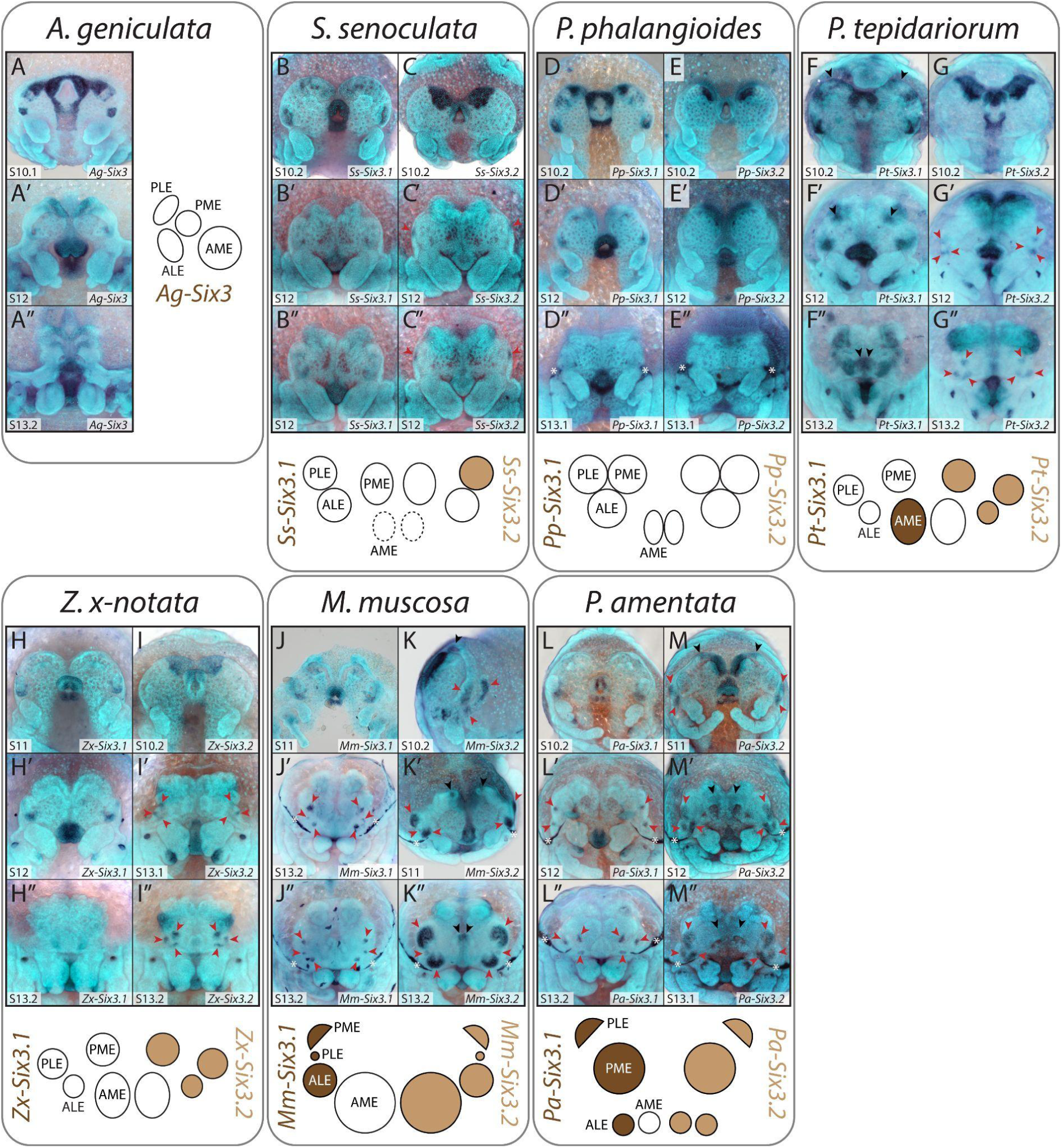
Expression of *Six3* orthologs in developing spider embryos. Two copies of *Six3* were recovered from each species. *Six3.1* orthologs were expressed in the secondary eyes of the RTA clade species, and in the AMEs of *Parasteatoda tepidariorum*. Expression of *Six3.2* orthologs was detected in all secondary eye primordia, starting from stage 10.2-12, in all entelegyne species examined, plus in the PLEs of *Segestria senoculata*. Black arrowheads indicate principal eye primordia, red arrowheads indicate secondary eye primordia, and asterisks indicate artifactual staining of the developing cuticle.

Across the other species examined, the overall pattern of *Six3* ortholog expression was somewhat conserved between species, but these genes were the most variable of the RDGs included in this study. In both *A. geniculata* and *P. phalangioides* none of the *Six3* orthologs analysed were detected in any of the eye primordia (Figure 6A-A” and 6D-E”). In *S. senoculata*, *Ss-Six3.2* was not expressed in any eye primordia, but *Ss-Six3.2* was expressed in a region that might overlap with the developing PLEs at stage 12 (Figure 6C’ and 6C”).

In *Z. x-notata*, *Zx-Six3.2* is expressed in all secondary eye primordia by stage 13.1, but again we did not detect *Zx-Six3.1* expression in any developing eyes (Figure 6H-I”). In *M. muscosa* and *P. amentata*, both *Six3* copies are expressed in all secondary eye primordia, but the timing and size of the expression domains differs between them (Figure 6J-M”). *Mm-Six3.1* and *Pa-Six3.1* expression is only visible from stage 12 onwards and is restricted to a small region at the centre of each secondary eye primordium (Figure 6J-J” and 6L-L”). *Mm-Six3.2* and *Pa-Six3.2* expression is already visible at stage 10.2/11 in three distinct domains corresponding to each secondary eye (Figure 6K, 6K’ and 6M, red arrowheads). By stage 13.2, the domain of *Mm-Six3.2* in the ALEs and PMEs is much larger with stronger expression stronger than in the PLEs (Figure 6K”), and domain of *Pa-Six3.2* expression is much larger in the PMEs and PLEs in comparison to the ALEs (Figure 6M”). Moreover, the larger expression domains of *Mm-Six3.2* in the PMEs and ALEs appear to surround the small domains of expression seen for *Mm-Six3.1* (Figure 6J” and 6K”). Additionally, expression of *Mm-Six3.2* and *Pa-Six3.2* is apparent in the AMEs of both species, being restricted to a small dorsal area of the AMEs in *M. muscosa* (Figure 6K-K” and 6M-M”).

### Expression of *dachshund* orthologs

In *P. tepidariorum*, *dac* expression is restricted to the secondary eyes. As previously described by Schomburg et al. (2015), *Pt-dac1* is first expressed in the early (stage 10.2) secondary eye primordia and later restricted to the ALEs (Figure 7G-G”), while *Pt-dac2* is expressed in all secondary eye primordia from stage 12 (Figure 7H-H”).

**Figure 7.**
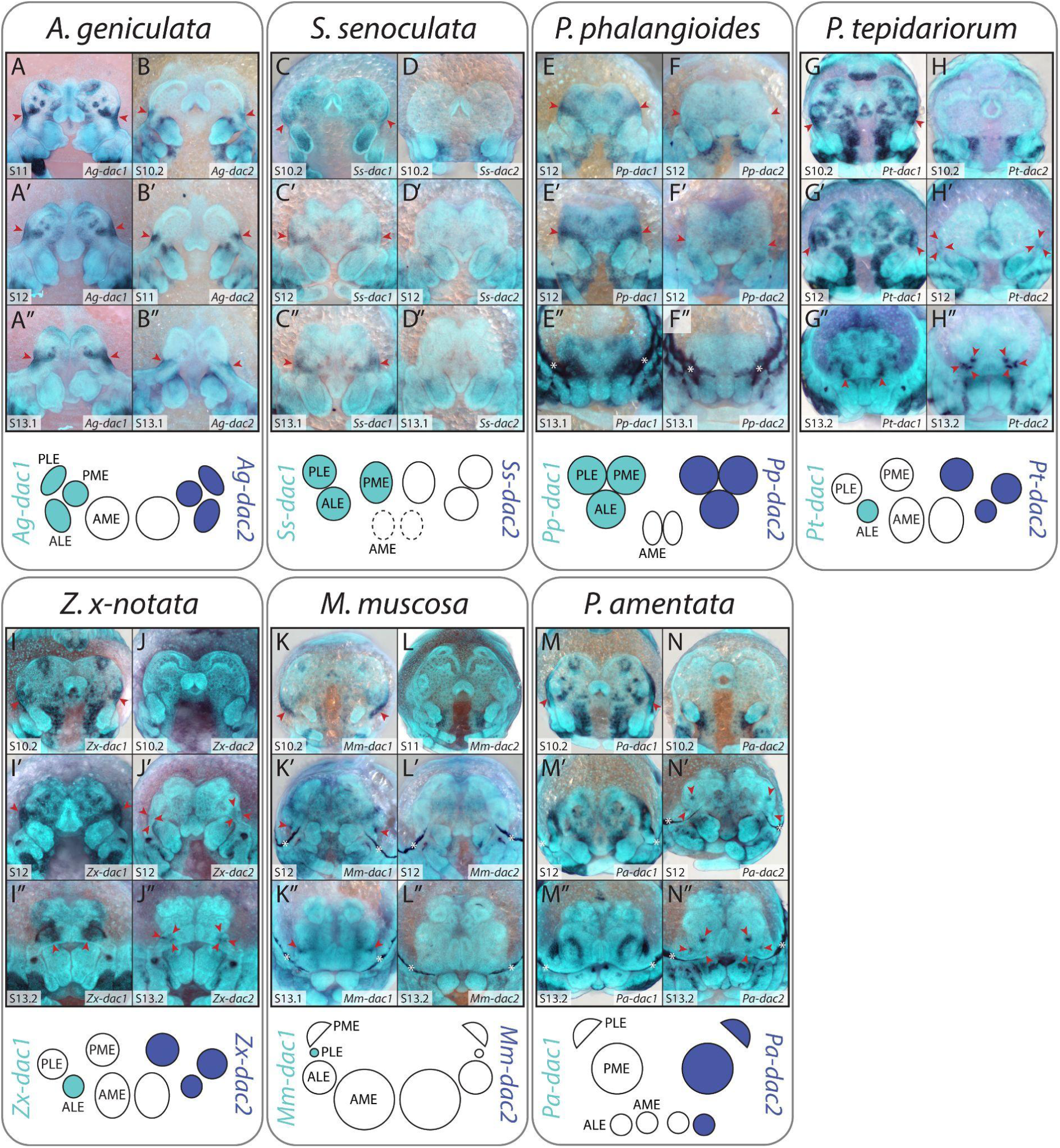
Expression of *dachshund* orthologs in developing spider embryos. Two copies of *dac* were recovered from each species. Orthologs of *dac1* were expressed in all secondary eyes in *A. geniculata, Segestria senoculata*, and *Pholcus phalangioides*, but only in the ALEs of *Parasteatoda tepidariorum* and *Zygiella x-notata* and in the PLEs of *Marpissa muscosa*. Expression of *dac2* was detected in all secondary eye primordia from stage 10.2-12 onwards in most species, but was absent in *M. muscosa* and *Acanthoscurria geniculata*. Black arrowheads indicate principal eye primordia, red arrowheads indicate secondary eye primordia, and asterisks indicate artifactual staining of the developing cuticle.

The early expression of *Pt-dac1* in the secondary eye primordia appears to be conserved with respect to all respective orthologs analysed (Figure 7A, 7C, 7E, 7G, 7I, 7K, 7M). However, there were differences in expression at later stages, with only *Zx-dac1* showing identical expression to *Pt-dac1* throughout (Figure 7I-I”). The early expression domains of *Ag-dac1, Pp-dac1*, and *Ss-dac1* in the secondary eye primordia do not appear to become restricted to any secondary eye pair, instead persisting as one contiguous domain even after the splitting of the individual eyes (using *so* expression as a marker) (Figure 7A-A”, 7C-C” and 7E-E”). In *P. amentata*, expression of *Pa-dac1* appears to partially surround the PLEs and PMEs by stage 12 (Figure 7M’ and 7M”), although these expression domains could partially overlap with the primordia of these secondary eyes. *Mm-dac1* expression is restricted to the PLEs from stage 12 onwards (Figure 7K’ and 7K”), rather than the ALEs. Expression of the *Pt-dac2* orthologs appears to be more variable among species. In *A. geniculata*, expression of *Ag-dac2* is first visible at stage 10.2 in the early secondary eye primordia. Like *Ag-dac1*, *Ag-dac2* expression does not split into distinct domains corresponding to individual eyes (Figure 7B-B”), but it appears to be more restricted in expression overall than the region of *Ag-dac1* expression (Figure 7A-B”). Likewise *Pp-dac2* is expressed in the secondary eye primordia from stage 12, but without distinct domains of expression corresponding to each secondary eye (Figure 7F-F”). Expression of *Zx-dac2* was also identical to *P. tepidariorum* (Figure 7J-J’’). Expression of *Pa-dac2* was detected in all secondary eye pairs, but the size of the expression domains did not reflect the size of the eyes as seen for *so* or *Six3.2* expression (Figure 7N-N’’). *Ss-dac2* and *Mm-dac2* expression appears to be absent from all eye primordia (Figure 7D-D” and 7L-L’’).

### Expression of *eyes absent* orthologs

Expression of *Pt-eya* has been previously reported from stage 10 in all eye primordia (Schomburg et al. 2015), similar to *Pt-so1* (Figure 8D-D”). This appears to be broadly conserved between all species surveyed (Figure 8A-F”), but with some variation in spatiotemporal expression patterns.

**Figure 8.**
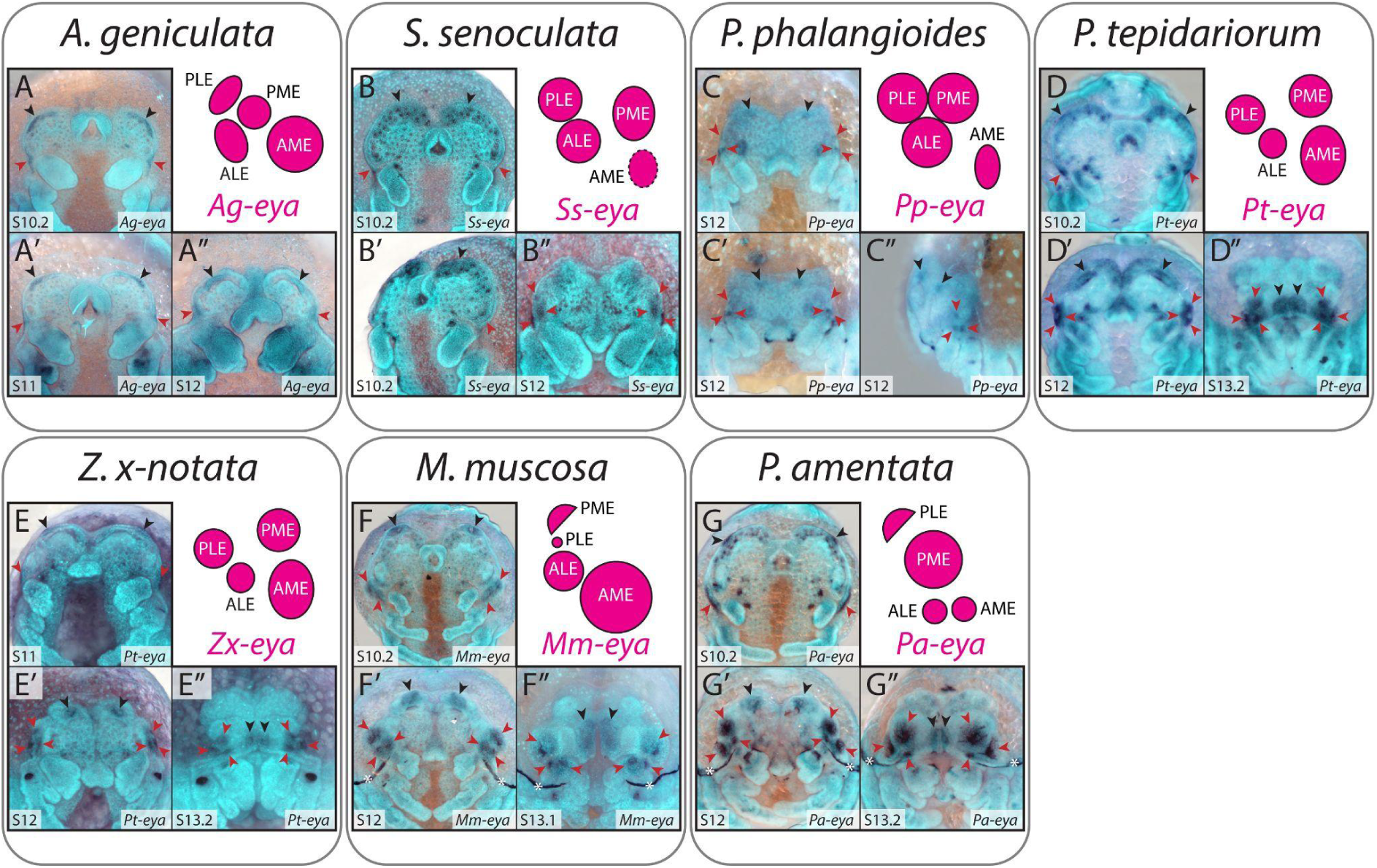
Expression of *eyes absent* orthologs in developing spider embryos. Only one copy of *eya* was recovered from each species. Expression of *eya* was detected in all eye primordia from stage 10.2 onwards, in all species examined. Black arrowheads indicate principal eye primordia, red arrowheads indicate secondary eye primordia, and asterisks indicate artifactual staining of the developing cuticle.

In *A. geniculata*, expression of *Ag-eya* in the secondary eye primordia does not seem to split into three pairs of discrete domains at stage 12 (Figure 8A-A”), similar to other *A. geniculata* RDGs. At stage 10.2, *Ss-eya* is expressed in a large region along the anterior edge of the head encompassing the principal eye primordia, as well as laterally around the secondary eye primordia, but by stage 12 expression is only visible in the latter (Figure 8B-B”). In *P. phalangioides*, the expression domains of *Pp-eya* do split into distinct primordia by stage 12 but appear to be smaller than those of *Pt-eya* and the expression of orthologs from other species (Figure 8C-C”).

*Zx-eya* is expressed in all eye pairs in an identical pattern to that of *Pt-eya* (Figure 8E-E”). In comparison to other *eya* orthologs at stage 12, *Mm-eya* has larger expression domains in the AMEs, ALEs and PMEs (Figure 8F-F”), with concentrated dots of *Mm-eya* expression within the ALEs and PMEs, and *Pa-eya* has larger expression domains in the PLEs and PMEs (Figure 8G-G”). However, expression of *Mm-eya* in the PLEs is restricted to a very small area (Figure 8F’ and 8F”) and expression of *Pa-eya* in the AMEs and ALEs is also reduced in size (Figure 8G’ and 8G”).

### Expression of *atonal* orthologs

*Pt-ato1* is expressed in all eye primordia as early as stage 10.2, with expression restricted to a few cells within each developing eye pair by stage 13.2 (Figure 9G-G”; Baudouin Gonzalez et al. 2022). Previously, we did not detect any expression of *Pt-ato2* in the eye primordia; however, in this study we observed expression of *Pt-ato2* in the primary eye primordia (Figure 9H-H”). This expression domain also appears to be restricted to a small area within the eye, but is more restricted than *Pt-ato1* (Figure 9G-H”).

**Figure 9.**
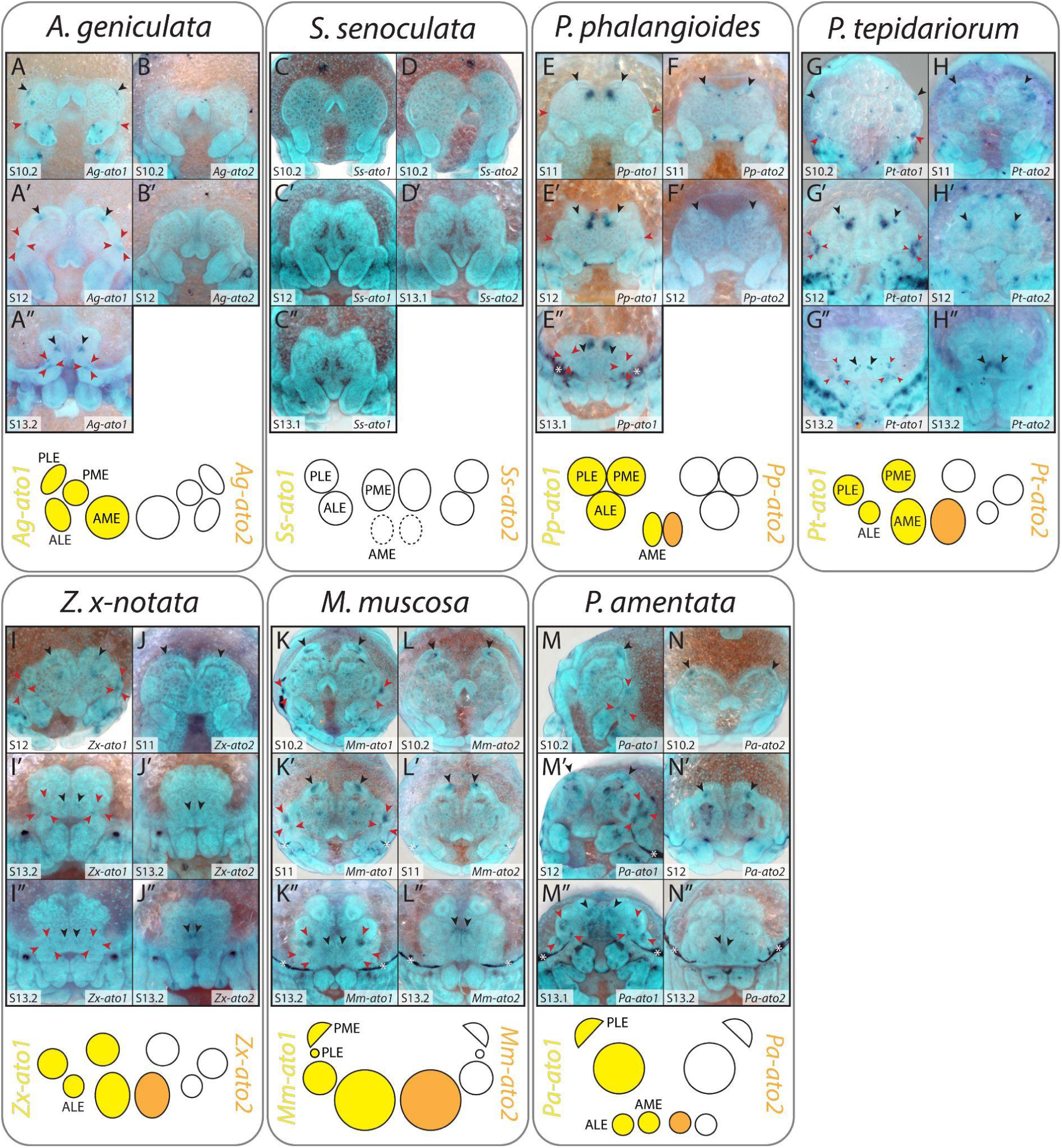
Expression of *atonal* orthologs in developing spider embryos. Two copies of *atonal* were recovered from each species. Expression of *ato1* orthologs was detected in all eye primordia from stage 10.2 onwards, with the exception of *Segestria senoculata*. Expression of *ato2* orthologs is restricted to the principal eyes in most species, but is absent in *S. senoculata* and *Acanthoscurria geniculata.* Black arrowheads indicate principal eye primordia, red arrowheads indicate secondary eye primordia, and asterisks indicate artifactual staining of the developing cuticle.

Expression of *ato* orthologs in the eye primordia is mostly conserved across species, with *Pt-ato1* orthologs being expressed in all eye primordia and *Pt-ato2* orthologs being expressed only in the primary eye primordia (Figure 9). However, there was some variation.

Firstly, we did not detect any expression of *Ag-ato2* in *A. geniculata* (Figure 9B and 9B’) or either copy of *ato* in *S. senoculata* (Figure 9C-D’) in the eye primordia. In *P. phalangioides*, expression of *Pp-ato1* in the secondary eye primordia did not split into three distinct domains at stage 12, and we could no longer detect expression in any eye primordia by stage 13.1 (Figure 9E-E”). The expression of *Zx-ato1* and *Zx-ato2* (Figure 9I-J”) were very similar to *P. tepidariorum*. In *M. muscosa*, we did not detect expression of *Mm-ato1* in the PLEs and, although expression is restricted to a smaller area within the other eye primordia, it does not seem to be as restricted as in *Pt-ato1* (Figure 9K-K”). The latter was also true of *Pa-ato1* expression in each eye primordia (Figure 9M-M”).

### Expression of *Pax6* orthologs

Previous work on *P. tepidariorum* did not detect any expression of the two *Pax6* orthologs in the eye primordia, with expression being restricted to the developing brain from stage 10.2 (Schomburg et al. 2015), which we have corroborated (Figure 10E-F”). *Pax6* ortholog expression was highly conserved in most species analysed, with expression being mainly restricted to the developing brain and absent from all eye primordia (Figure 10A-L”). However, both copies of *Pax6* appear to be expressed at lower levels in the non-neurogenic ectoderm along the edge of the developing head in *M. muscosa* and *P. amentata*, as well as *Ag-Pax6.2* in *A. geniculata* (Figure 10I-J” and 10B-B”).This *Ag-Pax6.2* expression could overlap with the eye primordia at stages 10.2 and 11, but not at stage 10.1 (Figure 10B-B”).

**Figure 10.**
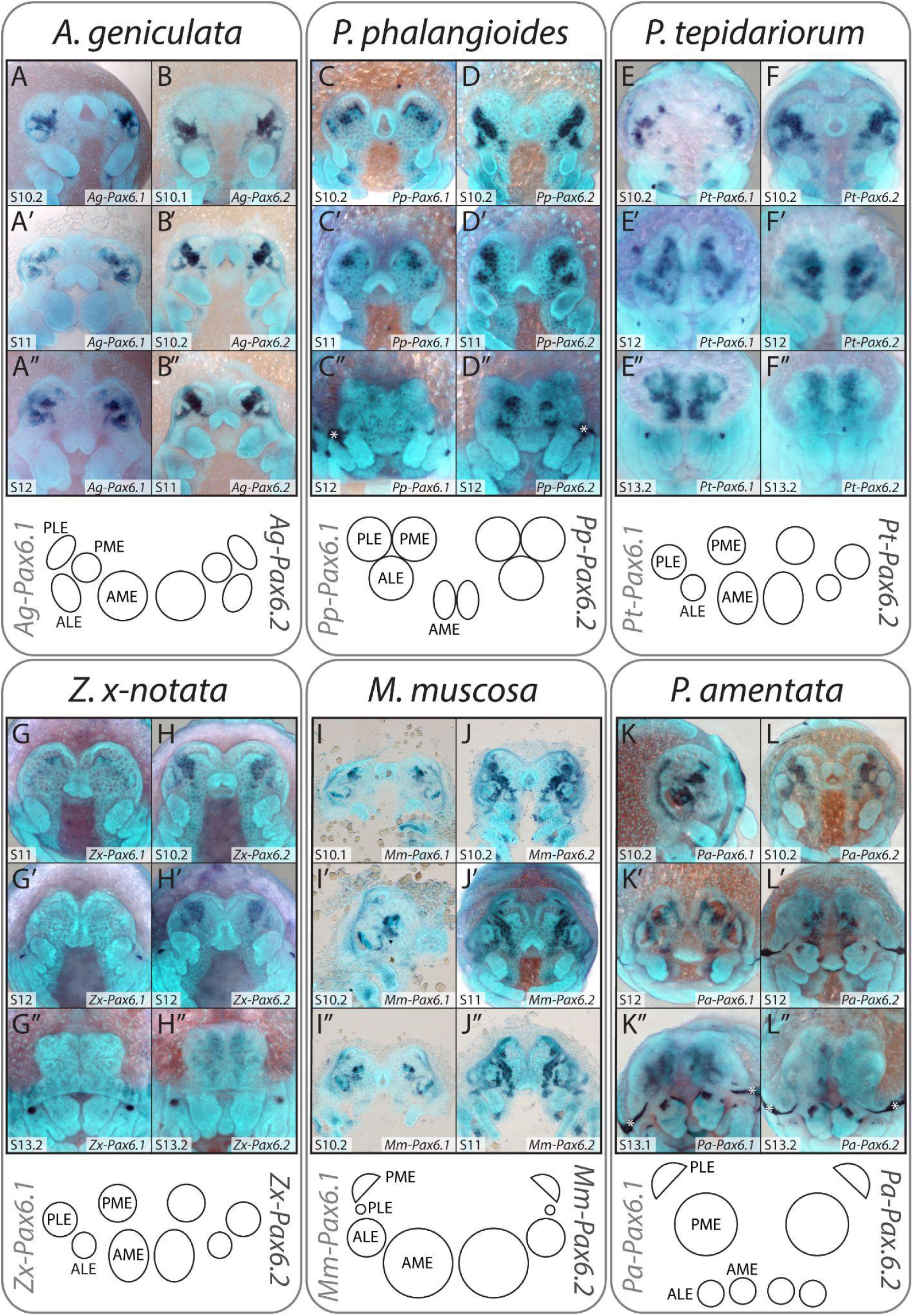
Expression of *Pax6* orthologs in developing spider embryos. Two copies of *Pax6* were recovered from each species, but no clear expression was detected in the eye primordia of any species. In some species, such as *Pardosa amentata*, *Pax6.1* expression appears to surround the secondary eye primordia, but in most, expression of *Pax6.2* is strongest and localises in the neurogenic ectoderm. Black arrowheads indicate position of principal eye primordia, red arrowheads indicate position of secondary eye primordia, and asterisks indicate artifactual staining of the developing cuticle.

In *Mm-Pax6.1* and *Pa-Pax6.1*, expression is restricted to the non-neurogenic ectoderm region between and proximal to the primary and secondary eye primordia at stage 10.2 (Figure 10I-I” and K). At stages 12 and 13.1, *Pa-Pax6.1* expression appears to partially surround the AMEs, ALEs and PMEs, which form gaps in the expression pattern (Figure 10K’ and 10K”).

*Mm-Pax6.2* and *Pa-Pax6.2* have similar expression patterns; there are gaps in expression that appear to correspond to the location of the secondary eye primordia (Figure 10 J-J” and 10L), and at stage 13.2, faint expression of *Pa-Pax6.2* may surround the ALEs, PLEs and PMEs (Figure 10L’ and 10L”). Additionally, *Mm-Pax6.2* expression in the non-neurogenic ectoderm may partially overlap with the primary eye primordia during stages 10.2-11 (Figure 10J-J”).

### Early expression of RDGs and eye initiation in P. tepidariorum

Leite et al. (2022) recently described stripes of early expression *of Pt-Pax6.1* and *Pt-Pax6.2* (from stage 6), which each appear at the anterior rim of the germ band. To establish whether these waves of expression could initiate eye development, we examined the early expression of *Pt-so1, Pt-eya*, *Pt-otd2,* and *Pt-Pax2.1*, which are the earliest known expressed RDGs from existing data for *P. tepidariorum* (Schomburg et al. 2015; Baudouin-Gonzalez et al. 2022; Janeschik et al. 2022; Leite et al. 2022). *Pt-eya* is expressed from early stage 8 around the dorsal periphery of the germ band and in the eye primordia by stage 10.2 (Figure 11A-B). Faint *Pt-so1* expression becomes visible at the perimeter of the head lobes at stage 8.2 (Figure 11C-D). This expression domain then splits, corresponding to the eye primordia by stage 10.1 (see Figure 4). By contrast, *Pt-otd2* expression is expressed in a stripe in the pre-cheliceral segment at stage 8.2 and only appears in the principal eye primordia from stage 10.1 onwards (Figure 11E-F). Double fluorescent ISH demonstrates that there is brief overlap of *Pt-Pax6.2 and Pt-otd2* expression in the developing brain, but not in the principal eye primordia (Figure 11G-I). *Pt-Pax2.1* is expressed in the undivided secondary eye primordia beginning at stage 10.1 (Figure 11J-L), slightly earlier than previously reported, and persists there until they separate.

**Figure 11.**
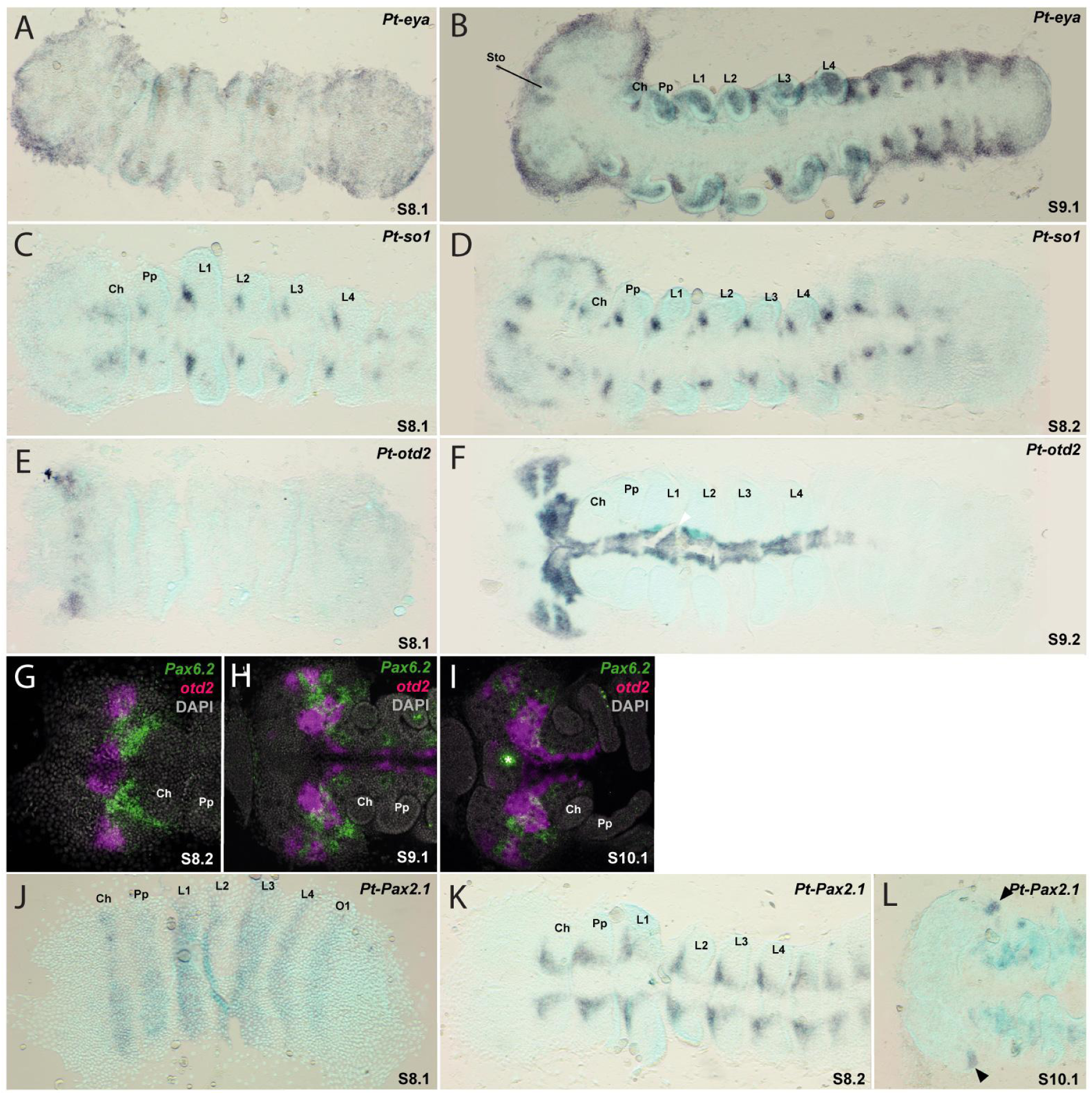
Early expression of *so1, eya, otd2,* and *Pax2.1* in *Parasteatoda tepidariorum.* Examination of stages 8.1-9.2 demonstrated that expression of *eyes absent* (A, B) and *sine oculis* (C, D) at the anterior rim of the head lobes begins at 8.1, earlier than previously appreciated. Although *otd2* expression is detectable in the head from stage 8.2, in the region of the developing eye primordia (E, F) it overlaps with *Pax6.2* expression only in the developing brain (G-I). Expression of *Pax2.1*, a marker for the secondary eye primordia, is not expressed during these early stages (J, K) but is detectable from stage 10.1 (L). Anterior is to the left in all images. Ch, cheliceral segment; L1-L4, walking leg segments 1-4; O1, opisthosomal segment 1; Pp, pedipalps.

### Selection analyses

Selection analyses using null and branch models in codeML revealed general trends in evolution of spider RDGs. First, we compared the dN/dS rate ratios (ω) between gene copies, labelling all spider lineages for each paralog as one group. All genes were highly conservative and no instances of positive selection, defined as ω>1, were identified in any group; however there were statistically significant (p < 0.05) differences in ω values for *ato* and *otd* duplications. In the case of *ato*, *ato2* (ω=0.066) exhibited significantly higher rates of non-synonymous change than *ato1* (ω=0.039). Similarly, among otd paralogs, *otd1* (ω=0.053) had a higher ω value compared to *otd2* (ω=0.017).

Next, we evaluated ω values for target spider lineages, based on their expression patterns and RDG repertoires. *ato* paralogs in *P. amentata* and *otd3* in *S. senoculata* were tested due to additional duplications, while *dac2* in *M. muscosa, Six3* paralogs in *M. muscosa* and *P. amentata*, and *so2* in *A. geniculata*, *M. muscosa*, *Z. x-notata*, *P. amentata*, *P. phalangioides*, *S. senoculata*, and *P. tepidariorum* were labelled as foregrounds due to their expression patterns. We also tested *otd* paralogs in *P. phalangioides* and *Pax6* in *A. geniculata.* Significant increase in ω values was observed for *dac2* in *M. muscosa* (ω1=0.07, ω0=0.028), *eyg* in *A. geniculata* (ω1=0.08, ω0=0.019), *so2* in *A. geniculata* (ω1=0.09, ω0=0.03), and *so2* in *P. phalangioides* (ω1=0.124, ω0=0.03). No additional synonymous substitutions were detected in *S. senoculata so2* compared to its nearest neighbour, *P. phalangioides so2*; thus, an exact value of ω is difficult to estimate, but positive selection is also likely to be acting on this branch. Significant decreases in ω, indicating lower selective pressure, was detected in *S. senoculata otd3* (ω1=0.011, ω0=0.036), *P. amentata Six3.2* (ω1=0.0001, ω0=0.016), *Z. x-notata so2* (ω1=0.006, ω0=0.031), and *P. tepidariorum so2* (ω1=0.01, ω0=0.031). The full results for each branch model are shown in Supplementary Table S3.

Branch-site models and Bayes empirical Bayes method indicated that two species had potentially positively selected codons, namely *M. muscosa* at sites 473 (glycine to glutamine) and 502 (glutamine to lysine) in *dac2,* and *A. geniculata* at sites 7 (alanine to asparagine), 209 (histidine to alanine), and 226 (glutamine to phenylalanine) in *so2.* The full summary of the branch-site tests is given in Supplementary Table S4.

## Discussion

These data provide the first detailed comparative insight to the developmental origins of visual system variation in spiders. Despite the strong conservation of RDG repertoires across the group, and more widely across arachnopulmonates, our data highlight several substantial differences in spatiotemporal expression patterns that could underpin key aspects of morphological, functional, and - ultimately - ecological diversity.

### Eye number

Although the vast majority of spiders have eight eyes, many taxa have lost one or more pairs at varying phylogenetic depths. This more commonly seems to affect the AMEs, presumably owing to their distinct developmental origins and regulatory networks, but one or more pairs of secondary eyes may also be lost. This phenomenon can be seen across large clades such as Dysderoidea or within individual lineages such as cavernicolous species of pholcids, nesticids, linyphiids, sparassids, and even lycosids (Jäger 2012; Mammola and Isaia 2017; Huber 2018). Interestingly, eye loss seems to occur particularly frequently in the plesiomorphic clade Synspermiata: likely independent losses appear in Caponiidae, Dysderoidea, Scytodoidea, and the Lost Trachea clade. The reduction and loss of the AMEs also appears commonly within Marronoidea (Wolff pers. comm.), but in neither this group nor the Synspermiata is there a clear ecological pattern as to which lineages reduce or lose their AMEs.

Despite the loss of the AMEs exhibited in *S. senoculata* likely occurring at the base of Dysderoidea, some 100-150 Ma (Shao and Li 2018), we detected early activation of the RDGs *otd*, *eya*, and *so* in the principal eye primordia up to stage 10/11. This is reminiscent of early eye development in eyeless subterranean species such as *Astyanax mexicanus*, wherein the eyes develop to a relatively advanced stage before apoptotic events in the lens trigger their arrest and degradation (see Jeffery 2009; Rétaux and Casane 2013; Wilkens and Strecker 2017 for reviews). However, such losses are usually much more recent evolutionary events, on the scale of hundreds of thousands of years rather than hundreds of millions (Sumner-Rooney 2018). The persistence of RDG expression in *S. senoculata*, although developmentally brief, hints at pleiotropic effects of these genes or upstream factors (Sumner-Rooney 2018). However, further detail is elusive at this stage, especially given the unknown identity of the overall regulators and initiators of spider eye determination (Schomburg et al. 2015; Baudouin-Gonzalez et al. 2022; Janeschik et al. 2022). One tempting lead is the presence of a duplication of *otd1* detected in *P. phalangioides* and *S. senoculata*, the two synspermiatan species sampled for this study. Selection analyses demonstrated that *otd1* is subject to greater positive selection than is *otd2* across all spiders, and the *otd1* duplicates in *P. phalangioides* and *S. senoculata* exhibit relatively long branches. However, this paralog is not expressed in the AMEs; instead, the *otd2* paralog is absent from *P. phalangioides* and still expressed in *S. senoculata*. Nevertheless, the possible correlation of this duplication with AME instability in Synspermiata might provide future insight and merits further investigation of *otd* function in eight- and six-eyed taxa. For example, tandem duplication and retention of *otd1* may reduce or remove stabilising selection on *otd2* in these lineages; indeed, *Ss-otd2* returned long branches and exhibited relaxed selection despite its expression persisting in the early AME primordia (Supplementary Table 3, Supplementary Figure 3). In cases where one or more pairs of secondary eyes are lost, the most parsimonious explanation would be failure of the secondary eye primordia to split into three distinct eye fields. Microstructural examination of the eyes of *Tetrablemma*, for example, reveal several merged eyes beneath shared lenses (Sumner-Rooney, unpublished observation).

### Eye size

Eye size is one of the most striking sources of variation in spider visual systems. It has direct functional implications for e.g. contrast sensitivity and achievable spatial resolution, which are most notably exploited by visual hunters such as salticids, deinopids, and lycosids (Blest and Land 1977; Land 1985; Winsor et al. 2023). In salticids, it was previously demonstrated that eye size is already established in juveniles and exhibits negative allometry (Goté et al. 2019). Our results demonstrate that, in salticids and lycosids, these differences in eye size are determined early in embryonic development; in *M. muscosa*, the enlargement of the AMEs is apparent in the *Mm-otd2* and *Mm-so1* expression domains as early as stage 10.2. Both *M. muscosa* and *P. amentata* exhibit enlargement in, and size variation between, the secondary eye pairs; a recent study by Chong et al. (2023), examining eye diameter across Araneae, found that this pattern is consistent across lineages that rely on visual cues for hunting. This enlargement is already visible in the overall size of the secondary eye primordium prior to its division into three distinct eye fields, and is particularly distinctive in *Mm-so2* and *Pa-so2* expression, compared to orthologs in other species. Samadi et al. (2015: Figure 4E) reported the same expression pattern of *so2* (as *six1a*) in *C. salei*, another visual hunter belonging to the RTA clade. The division of the primordium is also distinctly uneven in these species, such that the adult eye size variation is immediately apparent. This might reduce the need for unequal rates of cell proliferation in the resultant eye fields, but the control of field splitting remains unknown. In vertebrates, *hh* contributes to the division of the optic vesicle; while Baudouin-Gonzalez et al. (2022) did not report *hh* expression within the secondary eye primordia of *P. tepidariorum*, a very recent study discovered a previously unidentified second copy of *hh* in *P. tepidariorum, P. phalangioides, P. amentata*, and the mygalomorph *Ischnothele caudata* (Medina-Jiménez et al. submitted). This copy is expressed not only in the secondary eye primordia of these species, but in the principal eye primordia of (at least) *P. amentata*, *P. phalangioides*, and *I. caudata*. In *P. amentata*, *hh2* expression domains reflect size differences in the secondary eyes at stage 12, in line with our observations for *Pa-so1*, *Pa-eya*, and *Pa-Six3.2*. Whether this contributes to field splitting remains unclear; expression of *hh* in the vertebrate optic vesicle occurs at the centre and drives bilateral division by suppression between the eyes (e.g. Cardozo et al. 2014).

The determination of size could result from the activation of RDG expression in more cells, from increased cell proliferation within these regions, or both. Although the upstream factors responsible for the initial activation of RDGs in spiders is still unclear, Baudouin-Gonzalez et al. (2022) proposed that Wnt signalling restricts the eye field; we might therefore expect reduced Wnt expression in the surrounding head region to contribute to enlarged eye primordia.

Eye size is not reflected by the expression patterns of all the RDGs studied here. Regions of *atonal, dachshund,* and *Six3.1* expression appear to be restricted to certain parts of the developing eyes (see below) and therefore do not correspond so closely to overall size, a pattern also observed in *C. salei* (Samadi et al. 2015). Curiously, no RDG expression patterns in *P. phalangioides* reflected the difference between principal and secondary eye size exhibited in adults, which have small AMEs and larger secondary eyes of equal size (Chong et al. 2023). However, this pattern is less consistent among pholcids than lycosids or salticids, and it may be established later in development or in the early phases of post-developmental growth.

### Eye position

The arrangement of the eyes on the cephalothorax also varies substantially across Araneae (Morehouse et al. 2017). Common configurations include ‘halo’ arrangements, with two rows of eyes encircling the crown of the head (e.g. Araneidae), tight medial clusters or triads (e.g. Mygalomorphae), or a wider anterior-posterior distribution with frontal concentration of two or more eye pairs (e.g. Salticidae) (Figure 1). While the location of the AMEs is relatively consistent across large phylogenetic distances, the versatile placement of the secondary eye pairs facilitates this diversity. Our results suggest that the timing of the division of the secondary eye primordium could contribute to the eventual position of these eyes. In both *M. muscosa* and *P. amentata*, this division occurs earlier, around stage 10.2/11 as demonstrated by *eya*, *so1*, and *Six3.2* expression patterns (Figures 4-6). As a result, the PLEs are determined before head closure begins (Mittmann and Wolff 2012) and do not appear to migrate anteriorly as this process occurs. By contrast, the division of the secondary eye primordium was not reflected in *dac* expression patterns by stage 13.2 in *A. geniculata, S. senoculata*, or *P. phalangioides*, all of which exhibit clustered or triad arrangements with the secondary eyes being tight clustered and antero-medially located. *P. tepidariorum* and *Z. x-notata* sit somewhere between these two extremes, both in terms of timing and the eventual position of the secondary eyes.

### Eye identity, function, and regionalisation

Of course, eyes do not only vary in their size and location. They perform diverse functions, which can diverge both between lineages and between eye pairs, and they are often non-uniform in their structure.

The identity of the four eye pairs is apparently determined by a combinatorial code of RDG and potentially other genes, as already proposed by both Schomburg et al. (2015) and Samadi et al. (2015). From these two species, it is clear that this code varies between taxa, which we confirm here (Figure 12). The deep evolutionary distinction between the principal and secondary eyes is evidenced by their separate primordial origins and broadly conserved differences in the expression of e.g. *otd, dac, ato2,* and *Six3* orthologs, several of which are also reflected in the development of the ocelli and compound eyes of *D. melanogaster* (Friedrich 2003). Among the secondary eyes there is more variation between individual taxa as to which genes are expressed in the ALEs, PMEs, and PLEs specifically, including in *dac* and *Six3* expression. In *P. amentata* and previous descriptions of *C. salei* by Samadi et al. (2015), *dac2* expression surrounds the secondary eye primordia, rather than being present within the developing vesicles themselves.

**Figure 12.**
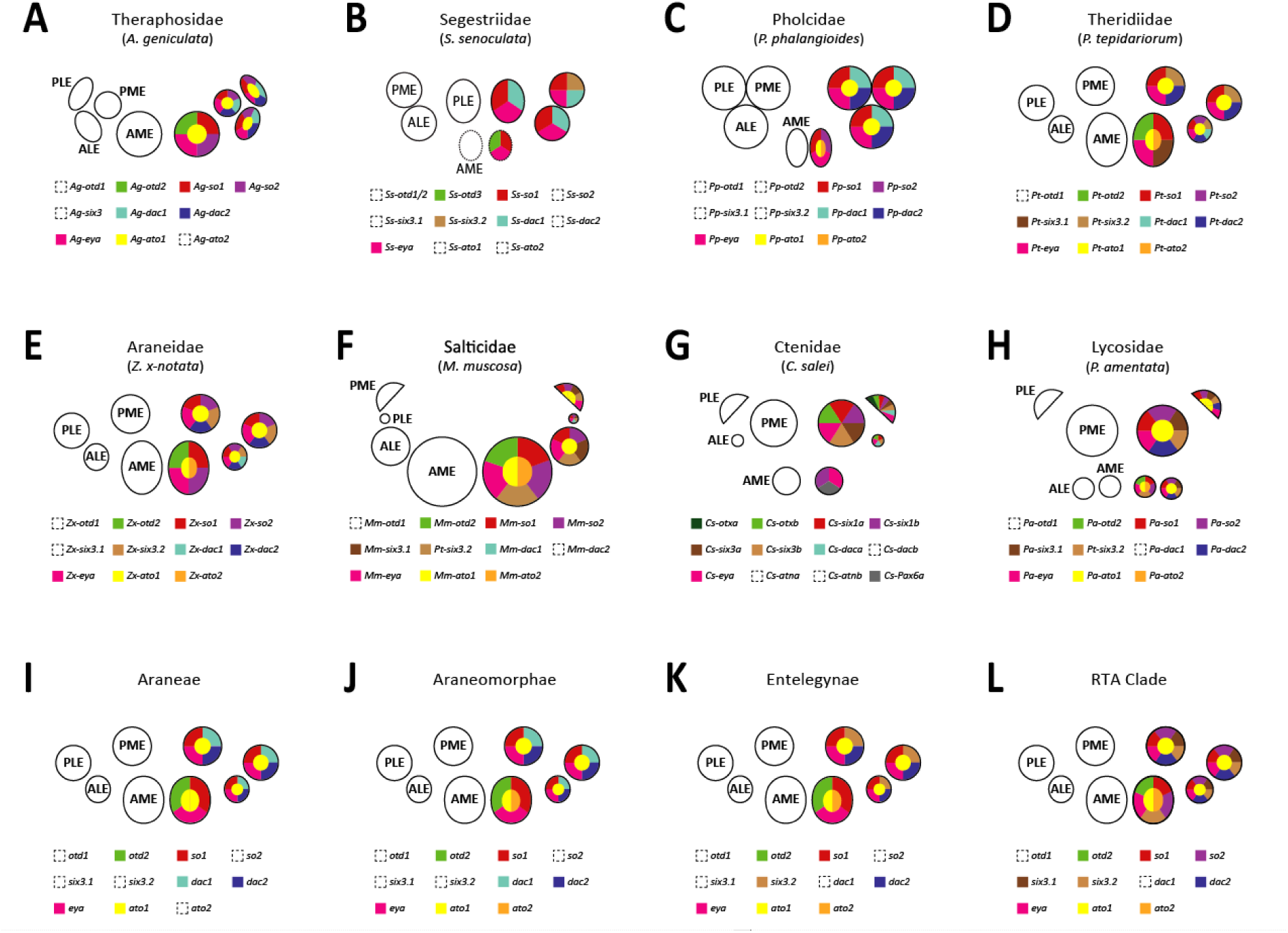
Summary of RDG expression across spiders. The combinations of RDGs expressed in each eye pair demonstrate aspects of both consistency and variation across the eight families studied to date (A-H). Phylogenetic patterns, such as the expression of *dac1* orthologs in all secondary eyes in the more plesiomorphic, non-entelegyne spiders, provide insight to the likely RDG expression in the ancestors of major spider clades (I-L).

The spatial resolution of our gene expression data is sufficient to identify cases of restricted expression within or surrounding the developing eyes. For example, *atonal* expression is restricted to small numbers of cells within the eyes; in *P. tepidariorum*, two distinct dots of *Pt-ato1* expression are clearly visible (Figure 9A’’). In other taxa such as *D. melanogaster*, *atonal* is regulates differentiation of photoreceptor cells (Jarman et al. 1994); Homann (1971) described the differentiation of the pigment and photoreceptor cells in two separate hemispheres of the retina, which corresponds to the observed expression patterns of *ato1*. The expression of *glass*, which acts downstream of *atonal* to trigger photoreceptor differentiation in *Drosophila* (Friedrich 2006), was also recently detected in the central part of the eyes of *P. tepidariorum* by Medina-Jiménez et al. (submitted), lending strong support to a conserved role for both genes in spiders. In *P. amentata, M. muscosa,* and *C. salei* (Samadi et al. 2015), regions of *so2* and *Six3.2* expression appear to be restricted to the distal edges of the eyes, while *Six3.1* is expressed in only a small central part of the secondary eyes. Whether these patterns are particular to the RTA clade or if they are simply more easily visible due to the size of the eyes, is unclear.

Whether, and how, the described differences in RDG expression might link to the morphological and functional distinctions between eyes, and between species, remains purely speculative. However, there are several intriguing correlations that are worthy of further exploration. For example, the detection of *ato2* expression in only the AMEs, given known links between *atonal* and photoreceptor differentiation in other organisms, could be connected to the determination of everted versus inverted photoreceptors in this eye type. The absence of *dac2* expression in the secondary eyes of *M. muscosa* was a striking and unexpected result; one unusual feature of the Salticidae is that their secondary eyes lack a tapetum (Homann 1971). We also detected site-specific positive selection in *Mm-dac2*, indicating possible functional change as well as differential regulation and expression. Future study of the closely related Philodromidae, which apparently share this loss of the tapetum (Homann 1971), might shed further light on this possible correlation.

### Eye initiation and the role of Pax6

One mystery that persists is how eye development is initiated in spiders (Friedrich 2022). Our expression data for six new species spread across the spider phylogeny demonstrate that there is no clear evidence for *Pax6* expression within the developing eye primordia. Although Samadi et al. (2015) reported expression in the developing AMEs in *C. salei*, we did not detect unequivocal overlap between the eye primordia and *Pax6* expression domains in any of the species examined. Instead, our data indicate that *Pax6* expression is localised within the developing central nervous system and potentially regions that correspond to the eventual optic neuropils from stages 9-10 onwards. This is congruent with previous reports of *Pax6* expression in the forebrain of many other invertebrates and vertebrates (Hartmann et al. 2003; Morante et al. 2011; Bandín et al. 2014; López et al. 2020). We also observed *Pax6.1* expression apparently surrounding the secondary eyes in *P. amentata*, but this pattern was not clear across other species.

However, the detection of stripes of *Pax6.1* and *Pax6.2* expression at the anterior rim of stage 6-7 embryos in *P. tepidariorum* by Leite et al. (2022), combined with our detection of *eya* and *so1* expression as early as stage 8.2, provides possible insight to the initiation of the visual system. Although *Pax6* and the expression of other RDG genes never overlap, it is possible that the wave of *Pax6* expression at the anterior rim of the embryo at stage 6-7 triggers later RDG expression in this area shortly afterwards. In the developing head of the beetle *Tribolium castaneum*, *Pax6* orthologs *eyeless* and *twin of eyeless* are required for patterning the ocular segment. The insect ocular segment habors precursor cells to the primordia of a number of peripheral and internal components of the larval and adult head, notably including the adult compound eyes, the larval eyes, and the optic neuropils (Luan et al. 2014). Similarly, in *P. tepidariorum Pax6.1* and *Pax6.2* are expressed in stripes from stage 6 to 8 that moves away from the anterior rim of the germ band to more posterior positions in the precheliceral lobes (Leite et al. 2022). Thus, components of the spider visual system likely originate from within the same early embryonic field of *Pax6*-expressing cells as in other arthropods (Friedrich 2022). Recent findings by Janeschik et al. (2022) identified *Pax2* as a new marker for the secondary eye primordia. The authors suggested that *Pax2* might play a role in the initiation of the secondary eyes in some spiders, including *P. phalangioides*, replacing the role of *Pax6*. The possibility of some overlap between the expression domains of *Pax6.2* and *so* orthologs in *A. geniculata*, combined with positive selection in *Ag-eyg*, is particularly interesting in this context, as these authors did not detect the expression of any *Pax2* orthologs in the embryonic transcriptome of this species. The earliest expression of *Pt-Pax2.1* was detected at stage 10.1; this is later than *Pt-so1* activation in the secondary eye primordia at stage 9.2 (Baudouin-Gonzalez et al. 2022).

### Phylogenetic patterns

Besides correlations between RDG expression patterns and adult morphology, we also see phylogenetic signal in our data, with RDG expression differing between major clades. Patterns that unite the more plesiomorphic non-entelegyne species (*A. geniculata, P. phalangioides,* and *S. senoculata*), such as the expression of *dac1* orthologs, but not *Six3.2* orthologs, in all secondary eye pairs, may reflect the ancestral state of spider visual system development. Likewise, only *A. geniculata* lacked obvious *ato2* expression in the developing AMEs, suggesting that this is specific to araneomorphs. *Ag-so2* exhibited significantly higher ω than its araneomorph orthologs, and was expressed in all eye primordia; while this corresponds to *so2* expression patterns in the more derived araneomorphs, it stands in contrast to its closest relatives in Synspermiata, where *so2* expression is restricted to the principal eye primordia. By contrast, there is also evidence for derived RDG expression patterns in specific clades and lineages: *M. muscosa* and *P. amentata* represent the highly derived RTA clade and share *Six3.1* expression in all secondary eye pairs, while both araneoids, *P. tepidariorum* and *Z. x-notata*, exhibited *dac1* expression restricted to the ALEs only.

### Is there a role for gene duplication?

Gene duplication can provide the opportunity for functional diversification via sub- or neofunctionalisation of duplicates (Ohno 1970; Holland et al. 1994; Holland 2013). Large-scale duplication events, such as the WGD seen in arachnopulmonates, provide copies of entire regulatory networks (Schwager et al. 2017; Leite et al. 2018, Aase-Remedios et al. 2023). Within spider visual systems, this could offer a route to divergence between the two eye types, or between individual pairs of secondary eyes (Samadi et al. 2015; Schomburg et al. 2015; Baudouin-Gonzalez et al. 2022). Although the potential contribution of WGD to spider diversification is clear, identifying specific instances of retained ohnologs (paralogs generated from WGDs) that have functionally diverged can be challenging. Here, there are some clear indications of such divergence that could support a role of WGD in visual system diversification. Ohnolog pairs that are both expressed in the developing visual system, but with distinct expression patterns, include *so, ato, dac,* and *Six3.* Of these, *Six3* expression most clearly shows signs of subfunctionalisation: in *M. muscosa, P. amentata*, and *C. salei,* the expression domains of *Six 3.1* and *Six3.2* appear to be mutually exclusive, with *Six3.2* expression surrounding that of *Six3.1* (Samadi et al. 2015). It is unclear whether this subfunctionalisation is specific to the RTA clade, or if it is more easily visible in the RTA clade due to their enlarged secondary eyes. It is curious, given this possible novel specialisation, that *Pa-Sx3.2* exhibited lower selection pressure than surrounding orthologs. Subfunctionalisation between *ato1* and *ato2* is also suggested by the typical expression of the former in all four eye pairs, but the restriction of the latter to the AMEs (in araneomorphs only, potentially highlighting a case of clade-specific subfunctionalisation). In addition to evidence from expression patterns, selection analyses also supported greater positive selection on *ato2* than *ato1*, and on *otd1* than *otd2*, indicating different selective regimes following duplication in these ohnologs.

## Conclusions

Spiders have exploited the modular nature of their visual systems to occupy a wide variety of ecological niches and morphospaces. We demonstrate that this diversity is underpinned by a highly conserved repertoire of RDGs, but that the spatial and temporal expression patterns of these genes, in many cases, reflect aspects of morphological and functional diversity in adults. We identify candidate genes and mechanisms involved in the determination of eye size, number, and position, as well as phylogenetic patterns and potential correlations to eye type and function. We find evidence for a contribution of gene duplication, through divergent expression patterns and asymmetrical evolution of paralogs, with potential implications for eye number, identity, and regionalisation.

## Author contributions

The project was conceived by LSR, APM, and LBG. Funding was obtained by LSR and APM. ISH was performed by LBG and AH. Transcriptome assembly, gene and transcript identification, and phylogenetic analysis were performed by LBG and LSR with assistance from SA, DJL, and VT. AS performed RNA extractions, library preparation, and culture maintenance. Specimens and bioinformatic resources for *M. muscosa* and *A. geniculata* were provided by POMS and MP. VT performed selection analyses with supervision by CK. AP and LSR reconstructed synchrotron scans. LSR and LBG wrote the first draft of the manuscript and produced figures with contributions from APM, AP, and AH. All authors edited and approved the manuscript.

## Acknowledgements

This work was funded by the Leverhulme Trust (RPG-2020-237), the John Fell Fund (University of Oxford, LS0005632), and Deutsche Forschungsgemeinschaft Emmy Noether programme (SU 1336/1-1). We acknowledge the Paul Scherrer Institut, Villigen, Switzerland for provision of synchrotron radiation beamtime at the TOMCAT beamline X02DA of the SLS (proposals 20210266 and 20190502) and warmly thank C.M. Schlepütz for assistance. We gratefully acknowledge Ivo Andrews, Laura Ashby, Grace Blakeley, Mark Carnall, Jack Matthews, Rochelle Meah, Vanessa Moore, and Chris Ward for their help collecting spiders, Natascha Turetzek for access to the assembled *P. phalangioides* transcriptome, Daniela Rößler and Gabriele Uhl for access to *M. muscosa* embryos and comments on the manuscript, and Sam J. England for photographs of *M. muscosa, P. amentata, P. phalangioides, S. senoculata,* and *Z. x-notata* used in Figure 1.

## Supplementary materials

See attached zip file, Baudouin-Gonzalez_etal_SupplementaryMaterials.zip.

## References

1. Akiyama-Oda, Y., and H. Oda. 2016. Multi-color FISH facilitates analysis of cell-type diversification and developmental gene regulation in the *Parasteatoda* spider embryo. Dev. Growth Differ. 58:215–224. John Wiley & Sons, Ltd.

2. Andrews, S. 2010.FastQC: a quality control tool for high throughput sequence data.

3. Arendt, D. 2003. Evolution of eyes and photoreceptor cell types. Int. J. Dev. Biol. 47:563–571.

4. Bandín, S., R. Morona, J. M. López, N. Moreno, and A. González. 2014. Immunohistochemical analysis of Pax6 and Pax7 expression in the CNS of adult *Xenopus laevis*. J. Chem. Neuroanat. 57–58:24–41.

5. Baudouin-Gonzalez, L., A. Harper, A. P. McGregor, and L. Sumner-Rooney. 2022. Regulation of eye determination and regionalization in the spider *Parasteatoda tepidariorum*. Cells 11:631. MDPI.

6. Baudouin-Gonzalez, L., A. Schoenauer, A. Harper, G. Blakeley, M. Seiter, S. Arif, L. Sumner-Rooney, S. Russell, P. P. Sharma, and A. P. McGregor. 2021. The evolution of Sox gene repertoires and regulation of segmentation in arachnids. Mol. Biol. Evol. 1–31.

7. Blest, A. D., and M. F. Land. 1977. The physiological optics of *Dinopis subrufus* L. Koch: a fish lens in a spider. Proc. R. Soc. Lond. - Biol. Sci. 196:197–222.

8. Buschbeck, E. K., and M. J. Bok (eds). 2023. Distributed vision: from simple sensors to sophisticated combination eyes. Springer.

9. Callaerts, P., J. Clements, C. Francis, and K. Hens. 2006. Pax6 and eye development in Arthropoda. Arthropod Struct. Dev. 35:379–391.

10. Camacho, C., G. Coulouris, V. Avagyan, N. Ma, J. Papadopoulos, K. Bealer, and T. Madden. 2009. BLAST+: architecture and applications. BMC Bioinformatics 10:421.

11. Capella-Gutiérrez, S., J. M. Silla-Martínez, and T. Gabaldón. 2009. trimAl: a tool for automated alignment trimming in large-scale phylogenetic analyses. Bioinformatics 25:1972–1973.

12. Cardozo, M. J., L. Sánchez-Arrones, Á. Sandonis, C. Sánchez-Camacho, G. Gestri, S. W. Wilson, I. Guerrero, and P. Bovolenta. 2014. Cdon acts as a Hedgehog decoy receptor during proximal-distal patterning of the optic vesicle. Nat. Commun. 5:4272.

13. Castresana, J. 2000. Selection of conserved blocks from multiple alignments for their use in phylogenetic analysis. Mol. Biol. Evol. 17:540–552.

14. Chen, Y.-C., and C. Desplan. 2020. Chapter Four - Gene regulatory networks during the development of the *Drosophila* visual system. Pp. 89–125 in I. S. Peter, ed. Current Topics in Developmental Biology. Academic Press.

15. Chong, K., A. Grahn, C. D. Perl, and L. Sumner-Rooney. 2023. Complex allometric relationships and ecological factors shape the development and evolution of eye size in the modular visual system of spiders. BioRXiv 2023.12.28.573503.

16. Dacke, M., D. E. Nilsson, E. J. Warrant, A. D. Blest, M. F. Land, and D. C. O’Caroll. 1999. Built-in polarizers form part of a compass organ in spiders. Nature 401:470–473.

17. Danecek, P., J. K. Bonfield, J. Liddle, J. Marshall, V. Ohan, M. O. Pollard, A. Whitwham, T. Keane, S. A. McCarthy, R. M. Davies, and H. Li. 2021. Twelve years of SAMtools and BCFtools. GigaScience 10:giab008.

18. Friedrich, M. 2022. Coming into clear sight at last: Ancestral and derived events during chelicerate visual system development. BioEssays 44:2200163.

19. Friedrich, M. 2006. Ancient mechanisms of visual sense organ development based on comparison of the gene networks controlling larval eye, ocellus, and compound eye specification in *Drosophila*. Arthropod Struct Dev. 35(4):357–78.

20. Friedrich, M. 2003. Evolution of insect eye development: First insights from fruit fly, grasshopper and flour beetle. Integr. Comp. Biol. 43:508–521.

21. Gaspar, P., I. Almudi, M. D. S. Nunes, and A. P. McGregor. 2019. Human eye conditions: insights from the fly eye. Hum. Genet. 138:973–991.

22. Gehring, W. J. 2014. The evolution of vision. Wiley Interdiscip. Rev. Dev. Biol. 3:1–40.

23. Gehring, Walter; Ikeo, K. 1999. Pax 6 and eye evolution. Trends Genet. 9525:371–377.

24. Goté, J. T., P. M. Butler, D. B. Zurek, E. K. Buschbeck, and N. I. Morehouse. 2019. Growing tiny eyes: How juvenile jumping spiders retain high visual performance in the face of size limitations and developmental constraints. Vision Res., doi: 10.1016/j.visres.2019.04.006.

25. Haas, B. J., A. Papanicolaou, M. Yassour, M. Grabherr, P. D. Blood, J. Bowden, M. B. Couger, D. Eccles, B. Li, M. Lieber, M. D. MacManes, M. Ott, J. Orvis, N. Pochet, F. Strozzi, N. Weeks, R. Westerman, T. William, C. N. Dewey, R. Henschel, R. D. LeDuc, N. Friedman, and A. Regev. 2013. De novo transcript sequence reconstruction from RNA-Seq: reference generation and analysis with Trinity.

26. Hartmann, B., P. N. Lee, Y. Y. Kang, S. Tomarev, H. G. de Couet, and P. Callaerts. 2003. Pax6 in the sepiolid squid *Euprymna scolopes*: evidence for a role in eye, sensory organ and brain development. Mech. Dev. 120:177–183.

27. Hoang, D. T., O. Chernomor, A. von Haeseler, B. Q. Minh, and L. S. Vinh. 2018. UFBoot2: Improving the ultrafast bootstrap approximation. Mol. Biol. Evol. 35:518–522.

28. Holland, L. Z. 2013. Evolution of new characters after whole genome duplications: Insights from amphioxus. Semin. Cell Dev. Biol. 24:101–109. Elsevier Ltd.

29. Holland, P. W. H., J. Garcia-Fernandez, N. A. Williams, and A. Sidow. 1994. Gene duplications and the origins of vertebrate development. Development 120:125–133.

30. Homann, H. 1971. Die Augen der Araneae. Zool. Morphol. Den Tiere 69:201–272.

31. Huber, B. A. 2018. Cave-dwelling pholcid spiders (Araneae, Pholcidae): A review. Subterr. Biol. 26:1–18.

32. Jäger, P. 2012. Revision of the genus *Sinopoda* Jäger, 1999 in Laos with discovery of the first eyeless huntsman spider species (Sparassidae: Heteropodinae). Zootaxa 57:37–57.

33. Jakob, E. M., S. M. Long, D. P. Harland, R. R. Jackson, A. Carey, M. E. Searles, A. H. Porter, C. Canavesi, and J. P. Rolland. 2018. Lateral eyes direct principal eyes as jumping spiders track objects. Curr. Biol. 28:R1092–R1093.

34. Janeschik, M., M. I. Schacht, F. Platten, and N. Turetzek. 2022. It takes two: discovery of spider Pax2 duplicates indicates prominent role in chelicerate central nervous system, eye, as well as external sense organ precursor formation and diversification after neo- and subfunctionalization. Front. Ecol. Evol. 10:1–20.

35. Jarman, A. P., E. H. Grell, L. Ackman, L. Y. Jan, and Y. N. Jan. 1994. *atonal* is the proneural gene for *Drosophila* photoreceptors. Nature. 369:398–400.

36. Jeffery, W. R. 2009. Regressive evolution in *Astyanax* cavefish. Annu. Rev. Genet. 43:25–47.

37. Kalyaanamoorthy, S., B. Q. Minh, T. K. F. Wong, A. Von Haeseler, and L. S. Jermiin. 2017. ModelFinder: Fast model selection for accurate phylogenetic estimates. Nat. Methods 14:587–589.

38. Katoh, K., and D. Standley. 2013. MAFFT multiple sequence alignment software version 7: improvements in performance and usability. Mol. Biol. Evol. 30:772–780.

39. Land, M. F. 1985. The Morphology and Optics of Spider Eyes. Pp. 53–78 in F. G. Barth, ed. Neurobiology of Arachnids. Springer-Verlag, Heidelberg.

40. Leite, D. J., A. Schönauer, G. Blakeley, A. Harper, H. Garcia-Castro, L. Baudouin-Gonzalez, R. Wang, N. Sarkis, A. Günther Nikola, V. Sai Poojitha Koka, N. J. Kenny, N. Turetzek, M. Pechmann, J. Solana, and A. P. McGregor. 2022. An atlas of spider development at single-cell resolution provides new insights into arthropod embryogenesis. bioRxiv, doi: 10.1101/2022.06.09.495456.

41. Li, W., and A. Godzik. 2006. Cd-hit: a fast program for clustering and comparing large sets of protein or nucleotide sequences. Bioinformatics 22:1658–1659.

42. López, J. M., R. Morona, N. Moreno, D. Lozano, S. Jiménez, and A. González. 2020. Pax6 expression highlights regional organization in the adult brain of lungfishes, the closest living relatives of land vertebrates. J. Comp. Neurol. 528:139–163.

43. Luan, Q., Q. Chen, and M. Friedrich. 2014. The Pax6 genes *eyeless* and *twin of eyeless* are required for global patterning of the ocular segment in the *Tribolium* embryo. Dev. Biol. 394:367–381.

44. Mammola, S., and M. Isaia. 2017. Spiders in caves. Proc. R. Soc. B 284:20170193.

45. Marone, F., and M. Stampanoni. 2012. Regridding reconstruction algorithm for real-time tomographic imaging. J. Synchrotron Radiat. 19:1029–1037. International Union of Crystallography.

46. Medina-Jiménez, B. I., G. Budd, M. Pechmann, and R. Janssen. Submitted. Single-cell sequencing reveals novel insights into spider eye development. Submitted in tandem with the current manuscript.

47. Mittmann, B., and C. Wolff. 2012. Embryonic development and staging of the cobweb spider *Parasteatoda tepidariorum* C. L. Koch, 1841 (syn.: *Achaearanea tepidariorum*; Araneomorphae; Theridiidae). Dev. Genes Evol. 222:189–216.

48. Morante, J., T. Erclik, and C. Desplan. 2011. Cell migration in *Drosophila* optic lobe neurons is controlled by eyeless/Pax6. Development 138:687–693.

49. Morehouse, N. 2020. Spider vision. Curr. Biol. 30:R975–R980. Elsevier.

50. Morehouse, N. I., E. K. Buschbeck, D. B. Zurek, M. Steck, and M. L. Porter. 2017. Molecular evolution of spider vision: new opportunities, familiar players. Biol. Bull. 233:21–38.

51. Nguyen, L. T., H. A. Schmidt, A. Von Haeseler, and B. Q. Minh. 2015. IQ-TREE: A fast and effective stochastic algorithm for estimating maximum-likelihood phylogenies. Mol. Biol. Evol. 32:268–274.

52. Nielsen, R., and Z. Yang. 1998. Likelihood models for detecting positively selected amino acid sites and applications to the HIV-1 envelope gene. Genetics 148:929–936.

53. Ohno, S. 1970. Evolution by Gene Duplication. Springer, New York.

54. Paganin, D., S. C. Mayo, T. E. Gureyev, P. R. Miller, and S. W. Wilkins. 2002. Simultaneous phase and amplitude extraction from a single defocused image of a homogeneous object. J. Microsc. 206:33–40.

55. Pechmann, M. 2020. Embryonic development and secondary axis induction in the Brazilian white knee tarantula *Acanthoscurria geniculata*, C. L. Koch, 1841 (Araneae; Mygalomorphae; Theraphosidae). Dev. Genes Evol. 230:75–94. Development Genes and Evolution.

56. Prpic, N.-M., M. Schoppmeier, and W. G. M. Damen. 2008. Gene silencing via embryonic RNAi in spider embryos. Cold Spring Harb. Protoc.

57. Quinlan, A. R., and I. M. Hall. 2010. BEDTools: a flexible suite of utilities for comparing genomic features. Bioinformatics 26:841–842.

58. Rétaux, S., and D. Casane. 2013. Evolution of eye development in the darkness of caves: adaptation, drift, or both? EvoDevo 4:26.

59. Samadi, L., A. Schmid, and B. J. Eriksson. 2015. Differential expression of retinal determination genes in the principal and secondary eyes of *Cupiennius salei* Keyserling (1877). EvoDevo 6:16. ???

60. Schindelin, J., I. Arganda-Carreras, E. Frise, V. Kaynig, M. Longair, T. Pietzsch, S. Preibisch, C. Rueden, S. Saalfeld, B. Schmid, J.-Y. Tinevez, D. J. White, V. Hartenstein, K. Eliceiri, P. Tomancak, and A. Cardona. 2012. Fiji - an Open Source platform for biological image analysis. Nat. Methods 9:676–682.

61. Schomburg, C., N. Turetzek, M. I. Schacht, J. Schneider, P. Kirfel, N.-M. Prpic, and N. Posnien. 2015. Molecular characterization and embryonic origin of the eyes in the common house spider *Parasteatoda tepidariorum*. EvoDevo 6:15.

62. Seppey, M., M. Manni, and E. M. Zdobnov. 2019. BUSCO: Assessing genome assembly and annotation completeness. Gene Predict. Methods Mol. Biol. 1962:227–245. Humana, New York, NY.

63. Shao, L., and S. Li. 2018. Early Cretaceous greenhouse pumped higher taxa diversification in spiders. Mol. Phylogenet. Evol. 127:146–155.

64. Song, L., and L. Florea. 2015. Rcorrector: Efficient and accurate error correction for Illumina RNA-seq reads. GigaScience 4:1–8. GigaScience.

65. Stampanoni, M., A. Groso, A. Isenegger, G. Mikuljan, Q. Chen, A. Bertrand, S. Henein, R. Betemps, U. Frommherz, P. Böhler, D. Meister, M. Lange, and R. Abela. 2006. Trends in synchrotron-based tomographic imaging: the SLS experience. Pp. 193–206 in Developments in X-Ray Tomography V. SPIE.

66. Sumner-Rooney, L. 2018. The kingdom of the blind: disentangling fundamental drivers in the evolution of eye loss. Integr. Comp. Biol. 58:372–385.

67. Suyama, M., D. Torrents, and P. Bork. 2006. PAL2NAL: robust conversion of protein sequence alignments into the corresponding codon alignments. Nucleic Acids Res. 34:W609–W612.

68. Wilkens, H., and U. Strecker. 2017. Mechnanisms of regressive evolution. Pp. 1–217 in Evolution in the Dark: Darwin’s Loss Without Selection.

69. Winsor, A. M., N. I. Morehouse, and E. M. Jakob. 2023. Distributed Vision in Spiders. Pp. 267–318 in Distributed Vision: From Simple Sensors to Sophisticated Combination Eyes. Springer.

70. Yang, Z. 1998. Likelihood ratio tests for detecting positive selection and application to primate lysozyme evolution. Mol. Biol. Evol. 15:568–573.

71. Yang, Z. 2007. PAML 4: Phylogenetic Analysis by Maximum Likelihood. Mol. Biol. Evol. 24:1586–1591.

72. Yang, Z., W. S. W. Wong, and R. Nielsen. 2005. Bayes empirical Bayes inference of amino acid sites under positive selection. Mol. Biol. Evol. 22:1107–1118.

73. Zhang, J., R. Nielsen, and Z. Yang. 2005. Evaluation of an improved branch-site likelihood method for detecting positive selection at the molecular level. Mol. Biol. Evol. 22:2472–2479.

74. Zurek, D. B., T. W. Cronin, L. A. Taylor, K. Byrne, M. L. G. Sullivan, and N. I. Morehouse. 2015. Spectral filtering enables trichromatic vision in colorful jumping spiders. Curr. Biol. 25:R403–R404.

