## Supplementary material for "Development and patterning of a highly versatile visual system in spiders": Baudouin-Gonzalez et al. supplementary materials.docx

**Supplementary materials	Baudouin-Gonzalez et al.**

*Supplementary Table 1: Primers used for the amplification of RDGs*

*Supplementary Table 2: Gene identifiers for RDGs*

*Supplementary Table 3: Estimated parameters of selection tests using branch models in codeML*

*Supplementary Table 4: Codon positions under positive selection detected by the branch-site model using codeML*

*Supplementary files 1-6: Nucleotide alignments of RDGs*

SF1 ato_nucl-align.fas

SF2 dachshund_nucl-align.fas

SF3 otd_nucl-align.fas

SF4 Pax6_nucl-align.fas

SF5 Six3_nucl-align.fas

SF6 so_nucl-align.fas

*Supplementary Figures 1-6: Phylogenetic trees of RDGs generated using IQ-TREE with ModelFinder.*

SF1 Phylogenetic relationships of ato_nucl-align.fas

SF2 Phylogenetic relationships of dachshund_nucl-align.fas

SF3 Phylogenetic relationships of otd_nucl-align.fas

SF4 Phylogenetic relationships of Pax6_nucl-align.fas

SF5 Phylogenetic relationships of Six3_nucl-align.fas

SF6 Phylogenetic relationships of so_nucl-align.fas
