## Supplementary material for "Development and patterning of a highly versatile visual system in spiders": Supplementary Figures 1-6.docx

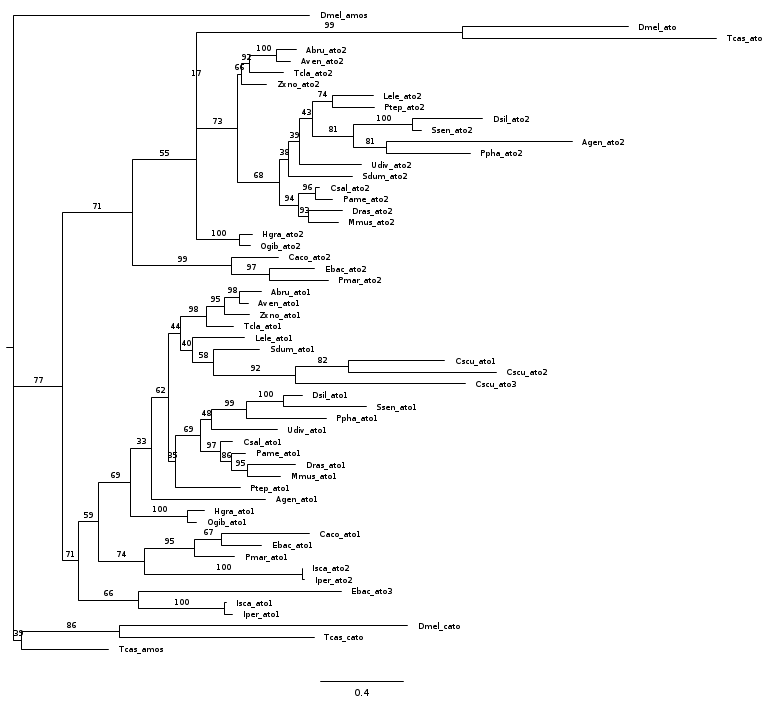

**SF1 Phylogeny for *ato* genes generated by IQ-TREE using ModelFinder**

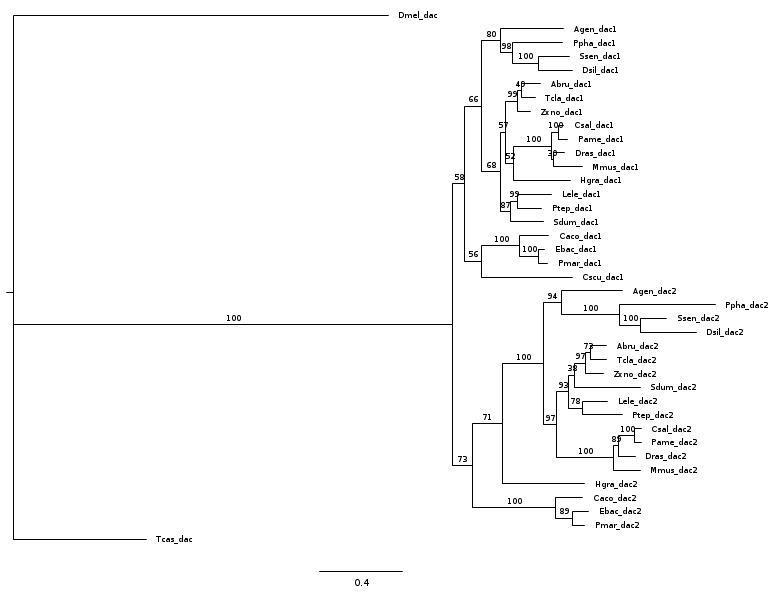

**SF2 Phylogeny for *dac* genes generated by IQ-TREE using ModelFinder**

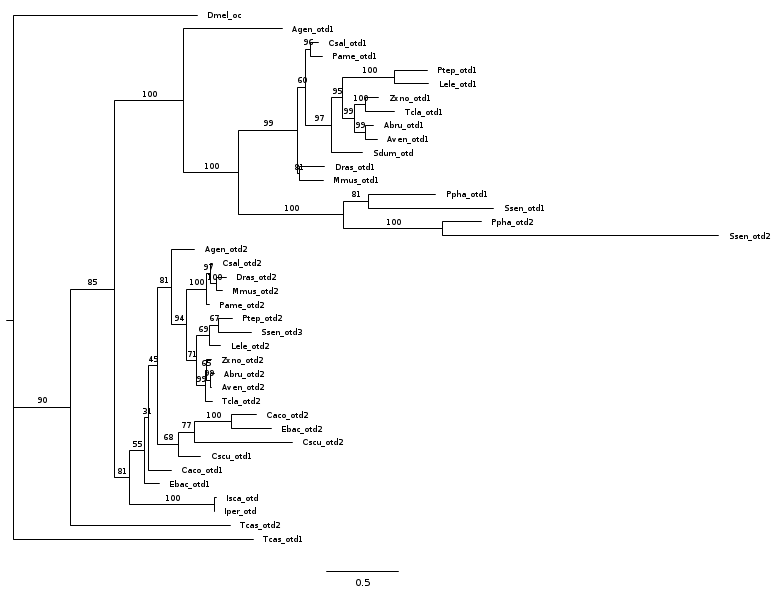

**SF3 Phylogeny for *otd* genes generated by IQ-TREE using ModelFinder**

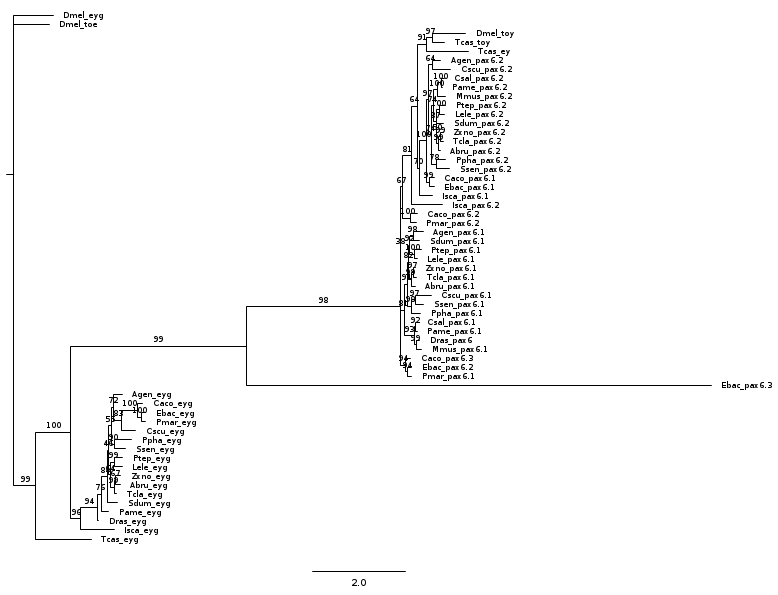

**SF4 Phylogeny for *Pax6* genes generated by IQ-TREE using ModelFinder**

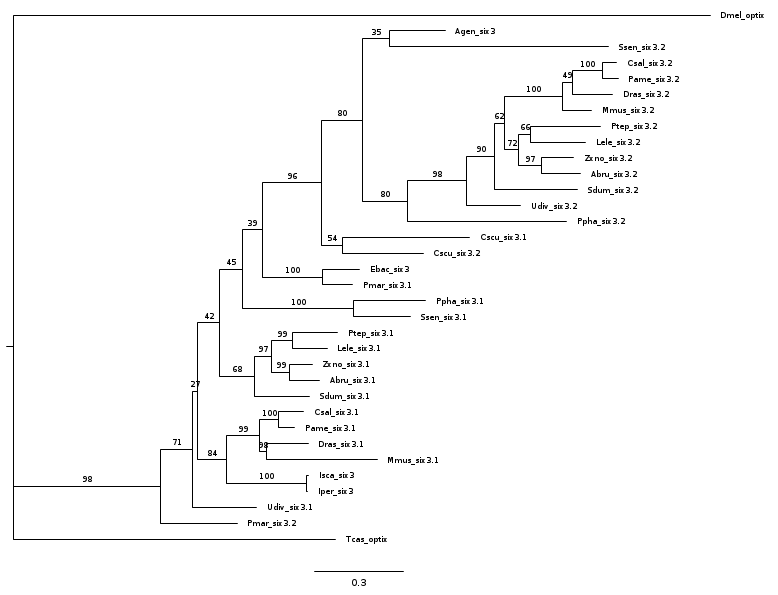

**SF5 Phylogeny for *Six3* genes generated by IQ-TREE using ModelFinder**

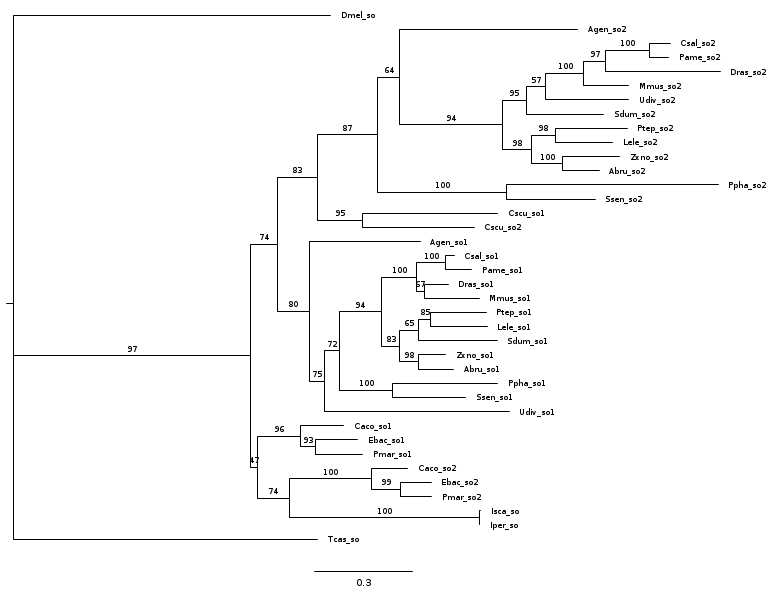

**SF6 Phylogeny for *so* genes generated by IQ-TREE using ModelFinder**
