## Supplementary material for "Development and patterning of a highly versatile visual system in spiders": Supplementary Tables 3-4.docx

**Table S3 Estimated parameters of selection tests using branch models in codeML**

| Model | Estimates of parameters | lnL | p |
| --- | --- | --- | --- |
| ***ato* paralogs** | | | |
| Null model M0 | *ω* = 0.051 | -12653.409 |  |
| **Alternative model 1 (*ato1* in spider lineages as foreground)** | ***ω*0 = 0.062,**  ***ω*1 = 0.039** | **-12648.198** | **0.001** |
| **Alternative model 2 (*ato2* in spider lineages as foreground)** | ***ω*0 = 0.042,**  ***ω*1 = 0.066** | **-12648.221** | **0.001** |
| Alternative model 3 (*Pardosa amentata ato1* and *ato2* as foreground) | *ω*0 = 0.051,  *ω*1 = 0.048 | -12653.397 | 0.877 |
| Alternative model 4 (*Pardosa amentata ato1* as foreground) | *ω*0 = 0.051,  *ω*1 = 0.048 | -12653.406 | 0.938 |
| Alternative model 5 (*Pardosa amentata ato2* as foreground) | *ω*0 = 0.051,  *ω*1 = 0.047 | -12653.400 | 0.893 |
| ***dac* paralogs** | | | |
| Null model M0 | *ω* = 0.029 | -28310.639 |  |
| Alternative model 1 (*dac1* in spider lineages as foreground) | *ω*0 = 0.029,  *ω*1 = 0.028 | -28310.605 | 0.794 |
| Alternative model 2 (*dac2* in spider lineages as foreground) | *ω*0 = 0.028,  *ω*1 = 0.03 | -28310.185 | 0.34 |
| **Alternative model 3 (*Marpissa muscosa dac2* as foreground)** | ***ω*0 = 0.028,**  ***ω*1 = 0.07** | **-28305.998** | **0.001** |
| ***otd* paralogs** | | | |
| Null model M0 | *ω* = 0.034 | -14087.424 |  |
| **Alternative model 1 (*otd1* in spider lineages as foreground)** | ***ω*0 = 0.018,**  ***ω*1 = 0.053** | **-14069.42** | **< 0.001** |
| **Alternative model 2 (*otd2* in spider lineages as foreground)** | ***ω*0 = 0.047,**  ***ω*1 = 0.017** | **-14072.574** | **< 0.001** |
| **Alternative model 3 (*Segestria senoculata otd2* as foreground)** | ***ω*0 = 0.036,**  ***ω*1 = 0.011** | **-14082.921** | **0.003** |
| Alternative model 4 (*Pholcus phalangioides otd1.1* and *otd1.2* as foreground) | *ω*0 = 0.033,  *ω*1 = 0.112 | -14086.578 | 0.193 |
| Alternative model 5 (*Pholcus phalangioides otd1.1* as foreground) | *ω*0 = 0.033,  *ω*1 = 0.167 | -14086.542 | 0.184 |
| Alternative model 6 (*Pholcus phalangioides otd1.2* as foreground) | *ω*0 = 0.028,  *ω*1 = 0.058 | -14087.332 | 0.668 |
| ***Pax6* paralogs** | | | |
| Null model M0 | *ω* = 0.02 | -26671.56 |  |
| Alternative model 1 (*eyg* in spider lineages as foreground) | *ω*0 = 0.02  *ω*1 = 0.02 | -26671.557 | 0.938 |
| Alternative model 2 (*Pax6.1* in spider lineages as foreground) | *ω*0 = 0.02  *ω*1 = 0.019 | -26671.259 | 0.438 |
| Alternative model 3 (*Pax6.2* in spider lineages as foreground) | *ω*0 = 0.02  *ω*1 = 0.019 | -26671.354 | 0.52 |
| **Alternative model 4 (*Acanthoscurria geniculata eyg* as foreground)** | ***ω*0 = 0.019**  ***ω*1 = 0.08** | **-26668.506** | **0.013** |
| Alternative model 5 (*Acanthoscurria geniculata Pax6.1* and *Pax6.2* as foreground) | *ω*0 = 0.019  *ω*1 = 0.027 | -26670.761 | 0.21 |
| Alternative model 6 (*Acanthoscurria geniculata Pax6.1* as foreground) | *ω*0 = 0.02  *ω*1 = 0.023 | -26671.488 | 0.7 |
| Alternative model 7 (*Acanthoscurria geniculata Pax6.2* as foreground) | *ω*0 = 0.02  *ω*1 = 0.017 | -26671.495 | 0.718 |
| ***Six3* paralogs** | | | |
| Null model M0 | *ω* = 0.016 | -12955.779 |  |
| Alternative model 1 (*Six3.1* in spider lineages as foreground) | *ω*0 = 0.019  *ω*1 = 0.013 | -12953.911 | 0.053 |
| Alternative model 2 (*Six3.2* in spider lineages as foreground) | *ω*0 = 0.018  *ω*1 = 0.013 | -12954.156 | 0.072 |
| Alternative model 3 (*Pardosa amentata Six3.1* and *Six3.2* as foreground) | *ω*0 = 0.016  *ω*1 = 0.01 | -12955.583 | 0.531 |
| Alternative model 4 (*Pardosa amentata Six3.1* as foreground) | *ω*0 = 0.015  *ω*1 = 0.028 | -12955.486 | 0.444 |
| **Alternative model 5 (*Pardosa amentata Six3.2* as foreground)** | ***ω*0 = 0.016**  ***ω*1 = 0.0001** | **-12953.799** | **0.047** |
| Alternative model 6 (*Marpissa muscosa Six3.1* and *Six3.2* as foreground) | *ω*0 = 0.016  *ω*1 = 0.01 | -12955.147 | 0.261 |
| Alternative model 7 (*Marpissa muscosa Six3.1* as foreground) | *ω*0 = 0.016  *ω*1 = 0.012 | -12955.442 | 0.412 |
| Alternative model 8 (*Marpissa muscosa Six3.2* as foreground) | *ω*0 = 0.016  *ω*1 = 0.007 | -12955.364 | 0.362 |
| ***so* paralogs** | | | |
| Null model M0 | *ω* = 0.03 | -16270.428 |  |
| Alternative model 1 (*so1* in spider lineages as foreground) | *ω*0 = 0.032  *ω*1 = 0.027 | -16269.616 | 0.203 |
| Alternative model 2 (*so2* in spider lineages as foreground) | *ω*0 = 0.029  *ω*1 = 0.032 | -16270.032 | 0.373 |
| Alternative model 3 (*so2* in species with uniform expression (*A. geniculata*,  *P. amentata*, *Z. x-notata*, *M. muscosa*) as foreground) | *ω*0 = 0.031  *ω*1 = 0.02 | -16268.985 | 0.089 |
| Alternative model 4 (*so2* in species with loss of expression (*P. phalangioides*,  *S. senoculata*, *P. tepidariorum*,) as foreground) | *ω*0 = 0.029  *ω*1 = 0.071 | -16268.867 | 0.077 |
| Alternative model 5 (*Pardosa amentata so2* as foreground) | *ω*0 = 0.031  *ω*1 = 0.009 | -16268.674 | 0.061 |
| Alternative model 6 (*Marpissa muscosa so2* as foreground) | *ω*0 = 0.03  *ω*1 = 0.044 | -16270.2 | 0.499 |
| **Alternative model 7 (*Acanthoscurria geniculata so2* as foreground)** | ***ω*0 = 0.03**  ***ω*1 = 0.09** | **-16268.468** | **0.048** |
| **Alternative model 8 (*Zygiella x-notata so2* as foreground)** | ***ω*0 = 0.031**  ***ω*1 = 0.006** | **-16264.536** | **< 0.001** |
| **Alternative model 9 (*Pholcus phalangioides so2* as foreground)** | ***ω*0 = 0.03**  ***ω*1 = 0.124** | **-16266.448** | **0.005** |
| **Alternative model 10 (*Segestria senoculata so2* as foreground)** | ***ω*0 = 0.03**  ***ω*1 = 999.000** | **-16266.367** | **0.004** |
| **Alternative model 11 (*Parasteatoda tepidariorum so2* as foreground)** | ***ω*0 = 0.031**  ***ω*1 = 0.01** | **-16267.755** | **0.021** |

**Table S4 Codon positions under positive selection detected by the branch-site model using codeML**

| Model | | lnL | p | BEB results  Pr(*ω*>1) > 95% |
| --- | --- | --- | --- | --- |
| ***ato* paralogs** | | | | |
| *Pardosa amentata ato1* and *ato2* as foreground | Null model (model A with *ω*2 = 1) | -12556.243 | 1 |  |
|  | Alternative model (Model A) | -12556.243 |  |  |
| *Pardosa amentata ato1* as foreground | Null model (model A with *ω*2 = 1) | -12556.243 | 1 |  |
|  | Alternative model (Model A) | -12556.243 |  |  |
| *Pardosa amentata ato2* as foreground | Null model (model A with *ω*2 = 1) | -12556.243 | 0.329 |  |
|  | Alternative model (Model A) | -12556.145 |  |  |
| ***dac* paralogs** | | | | |
| ***Marpissa muscosa dac2* as foreground** | **Null model (model A with *ω*2 = 1)** | **-28288.989** | **0.032** | **473 G → Q,**  **502 Q → K** |
|  | **Alternative model (Model A)** | **-28290.693** |  |  |
| ***otd* paralogs** | | | | |
| *Segestria senoculata otd2* as foreground | Null model (model A with *ω*2 = 1) | -14084.709 | 1 |  |
|  | Alternative model (Model A) | -14084.709 |  |  |
| *Pholcus phalangioides otd1.1* and *otd1.2* as foreground | Null model (model A with *ω*2 = 1) | -14062.481 | 0.235 |  |
|  | Alternative model (Model A) | -14062.220 |  |  |
| *Pholcus phalangioides otd1.1* as foreground | Null model (model A with *ω*2 = 1) | -14078.278 | 1 |  |
|  | Alternative model (Model A) | -14078.278 |  |  |
| *Pholcus phalangioides otd1.2* as foreground | Null model (model A with *ω*2 = 1) | -14078.465 | 0.076 |  |
|  | Alternative model (Model A) | -14077.439 |  |  |
| ***Pax6* paralogs** | | | | |
| *Acanthoscurria geniculata eyg* as foreground | Null model (model A with *ω*2 = 1) | -26671.559 | 1 |  |
|  | Alternative model (Model A) | -26671.559 |  |  |
| ***Acanthoscurria geniculata Pax6.1* and *Pax6.2* as foreground** | **Null model (model A with *ω*2 = 1)** | **-26671.559** | 0.003 |  |
|  | **Alternative model (Model A)** | **-26667.782** |  |  |
| *Acanthoscurria geniculata Pax6.1* as foreground | Null model (model A with *ω*2 = 1) | -26671.559 | 1 |  |
|  | Alternative model (Model A) | -26671.559 |  |  |
| *Acanthoscurria geniculata Pax6.2* as foreground | Null model (model A with *ω*2 = 1) | -26671.559 | 1 |  |
|  | Alternative model (Model A) | -26671.559 |  |  |
| ***Six3* paralogs** | | | | |
| *Marpissa muscosa Six3.1* and *Six3.2* as foreground | Null model (model A with *ω*2 = 1) | -12899.26 | 1 |  |
|  | Alternative model (Model A) | -12899.26 |  |  |
| *Marpissa muscosa Six3.1* as foreground | Null model (model A with *ω*2 = 1) | -12898.627 | 0.312 |  |
|  | Alternative model (Model A) | -12898.507 |  |  |
| *Marpissa muscosa Six3.2* as foreground | Null model (model A with *ω*2 = 1) | -12899.494 | 1 |  |
|  | Alternative model (Model A) | -12899.494 |  |  |
| *Pardosa amentata Six3.1* and *Six3.2* as foreground | Null model (model A with *ω*2 = 1) | -12899.494 | 1 |  |
|  | Alternative model (Model A) | -12899.494 |  |  |
| *Pardosa amentata Six3.1* as foreground | Null model (model A with *ω*2 = 1) | -12899.494 | 1 |  |
|  | Alternative model (Model A) | -12899.494 |  |  |
| *Pardosa amentata Six3.2* as foreground | Null model (model A with *ω*2 = 1) | -12899.494 | 1 |  |
|  | Alternative model (Model A) | -12899.494 |  |  |
| ***so* paralogs** | | | | |
| *Pardosa amentata so2* as foreground | Null model (model A with *ω*2 = 1) | -16226.887 | 1 |  |
|  | Alternative model (Model A) | -16226.887 |  |  |
| *Marpissa muscosa so2* as foreground) | Null model (model A with *ω*2 = 1) | -16226.887 | 1 |  |
|  | Alternative model (Model A) | -16226.887 |  |  |
| ***Acanthoscurria geniculata so2* as foreground** | **Null model (model A with *ω*2 = 1)** | **-16216.774** | **0.026** | **7 A → N,**  **209 H → A,**  **226 Q → F,** |
|  | **Alternative model (Model A)** | **-16214.877** |  |  |
| *Zygiella x-notata so2* as foreground | Null model (model A with *ω*2 = 1) | -16226.887 | 1 |  |
|  | Alternative model (Model A) | -16226.887 |  |  |
| *Pholcus phalangioides so2* as foreground | Null model (model A with *ω*2 = 1) | -16208.906 | 0.052 |  |
|  | Alternative model (Model A) | -16207.587 |  |  |
| *Segestria senoculata so2* as foreground | Null model (model A with *ω*2 = 1) | -16223.574 | 0.123 |  |
|  | Alternative model (Model A) | -16222.901 |  |  |
| *Parasteatoda tepidariorum so2* as foreground | Null model (model A with *ω*2 = 1) | -16226.887 | 1 |  |
|  | Alternative model (Model A) | -16226.887 |  |  |
